# Diet driven differences in host tolerance are linked to shifts in global gene expression in a common avian host-pathogen system

**DOI:** 10.1101/2024.08.07.607042

**Authors:** Erin L. Sauer, Carson Stacy, Weston Perrine, Ashley C. Love, Jeffrey A. Lewis, Sarah E. DuRant

## Abstract

As humans alter the landscape, wildlife have become increasingly dependent on anthropogenic resources, altering interactions between individuals and subsequently disease transmission dynamics. Further, nutritional quantity and quality greatly impact an individual host’s immune capacity and ability to mitigate damage caused by infectious disease. Thus, understanding the impact of dietary nutrition on immune function is critical for predicting disease severity and transmission as human activity both facilitates the dispersal of pathogens and alters dietary options for wildlife. Here, we use transcriptomics to explore the previously unstudied molecular mechanisms underpinning diet-driven differences in pathogen tolerance using a widespread avian bacterial pathogen, *Mycoplasma gallisepticum* (MG). MG is an ideal model for understanding the dietary drivers of disease as the human supplementation that wild birds commonly rely on, bird feeders, are also an important source for MG transmission. Significant diet-driven differences in the expression of many genes encoding immune response and translational machinery proteins are seen both in the absence of MG and during the recovery period. Prior to infection, protein-fed birds are more transcriptionally primed for infection than lipid-fed birds which translates to greater tolerance in protein-fed birds during the recovery period. Given the significant importance of human supplemented food in wildlife disease systems, the molecular mechanisms by which interactions between diet and infection emerge provide insight into the ecological and immunological consequences of human behavior and wildlife disease.

## Introduction

The availability, quality, and distribution of resources available to wildlife have drastically changed due to human activity and will continue to shift as humans further alter the landscape. Wildlife in human-dominated landscapes are becoming increasingly dependent on anthropogenic food sources, leading to changes in inter- and intra-specific interactions that increase disease transmission (1). Simultaneously, human activity has facilitated the dispersal of pathogens, leading to an increase in emerging infectious diseases that have caused catastrophic declines in biodiversity and pose a major threat to agriculture and human health (2). Defense against infectious disease requires energetically costly immune responses (3, 4). Nutritional resource quality and quantity can therefore shape both an individual host’s ability to cope with infection (3–6) and population-level disease dynamics (6–9). Understanding the impact of dietary nutrition on immune function and disease outcomes is thus critical for predicting disease dynamics as anthropogenic food sources become a greater proportion of wildlife diets and anthropogenic change alters wild dietary options (10, 11).

While both food quality and quantity greatly impact disease outcomes, most ecological research has focused on the availability of food, in part because it is difficult to disentangle the effects of caloric content (i.e. quantity) from nutrient content (i.e. quality;, 10). However, the nutritional content of food is also important in shaping disease outcomes and transmission (12). For example, micronutrient deficiencies in desert tortoises caused by anthropogenic habitat degradation subsequently led to immunodeficiency and severe disease driven population declines (13). Elucidating the individual immune components that are affected by nutritional quality is also difficult, as vertebrate immune systems are especially complex, relying on hundreds of molecular pathways (14). Thus, the impact of dietary nutritional quality on the molecular mechanisms of host responses to disease are especially under explored in wildlife-disease systems despite simultaneous increases in wildlife reliance on anthropogenic food sources and emerging diseases (15).

While the macronutritional content of food (fats, proteins, and carbohydrates) is critical to all aspects of health and cellular function in living organisms, including immune defense, its effects on cellular function are varied and complex (4, 5, 12). Amino acids are necessary for the synthesis of vital immune proteins such as cytokines and antibodies as well as the regulation and activation of the innate and adaptive immune system (4). Lipids also play a critical role in the immune system, particularly in modulating the expression of numerous inflammation genes (12, 16–18). Micronutrient content can also influence immune function, for example, various micronutrients alter inflammation pathways and T-cells expression in chickens (19, 20). However, all of these nutritional resources are also available to the pathogen and hosts commonly use self-induced anorexia during infection to restrict pathogen access to nutrients (21). Thus, restriction studies and self-induced anorexia complicate our ability to parse the effects of diet quality and quantity (caloric content) on immune function, especially in non-model systems.

Here, we explore the previously unstudied molecular mechanisms underpinning diet-driven differences in pathogen tolerance in a widespread avian bacterial pathogen, *Mycoplasma gallisepticum* (hereafter MG), which is both a conservation concern for wild birds and an economic problem for poultry farming. MG was detected shortly after the pathogen’s emergence in wild birds during the mid-1990’s causing large declines of house finch (*Haemorhous mexicanus*) populations in their introduced range of eastern North America and outbreaks in numerous other bird species (22–24). MG is spread through fomites and direct contact with infected birds, with bird feeders being important drivers of transmission in this system (25–29). Additionally, the eastern introduced house finch population is almost exclusively found in urban and suburban habitats where supplemental food from bird feeders make up a substantial portion of their diets (30). Because of the importance of bird feeders in this system, the type of diet birds are getting from feeders is also likely to play a role in disease dynamics making this an ideal system for understanding the dietary drivers of disease. Determining the currently unexplored effects of diet on molecular immune mechanisms improves our understanding of heterogeneity in disease outcomes among individuals and populations in this system.

To date there are no studies examining the molecular mechanisms that underpin heterogeneity in tolerance driven by ecological factors (e.g. diet) in the MG system. Nearly all transcriptomic studies in this system have focused on evolutionary questions (31–34). These studies have found that various pro-inflammatory pathways are typically downregulated in more tolerant house finch populations with endemic MG when compared to populations with little to no coevolutionary history (31, 33). Additionally, these studies focus on early and peak infection as this is known to be the stage where cytokine expression, and conjunctival inflammation, is greatest (31, 33, 34). However, gene expression prior to infection, which sets the stage for the immediate immunological response, and in the recovery period, a critical stage for predicting epidemic size, are also of interest (31, 35, 36). Thus, a better understanding of how ecological factors affect immune mechanisms both before and after pathogen exposure in wildlife is needed.

In prior experimental work on domestic canaries (*Serinus canaria domestica*) infected with MG, we found that a lipid rich diet led to prolonged pathology, but not differences in pathogen resistance, when compared to protein rich diets (Figure 1;, 37). This diet-driven difference in tolerance (defined as pathology per unit pathogen) should have impacts on MG survival and transmission, as both are heavily influenced by pathology (i.e. conjunctivitis intensity) and recovery time (36, 38–41). Here, we use transcriptomics to quantify gene expression in whole blood samples collected during the experiments described above prior to MG exposure and during the recovery period (14 & 21 days post exposure Figure 1A), where we found the greatest difference in tolerance (Figure 1B). We predict that prior to infection, expression of immune related genes will be influenced by diet. Further, we expect that during the recovery period immune genes and, specifically, inflammation process genes in tolerant protein-fed birds will be downregulated compared to lipid-fed birds. Finally, we predict that tolerance to MG will correspond to a smaller change in gene expression of genes related to the immune response.

**Figure 1.**
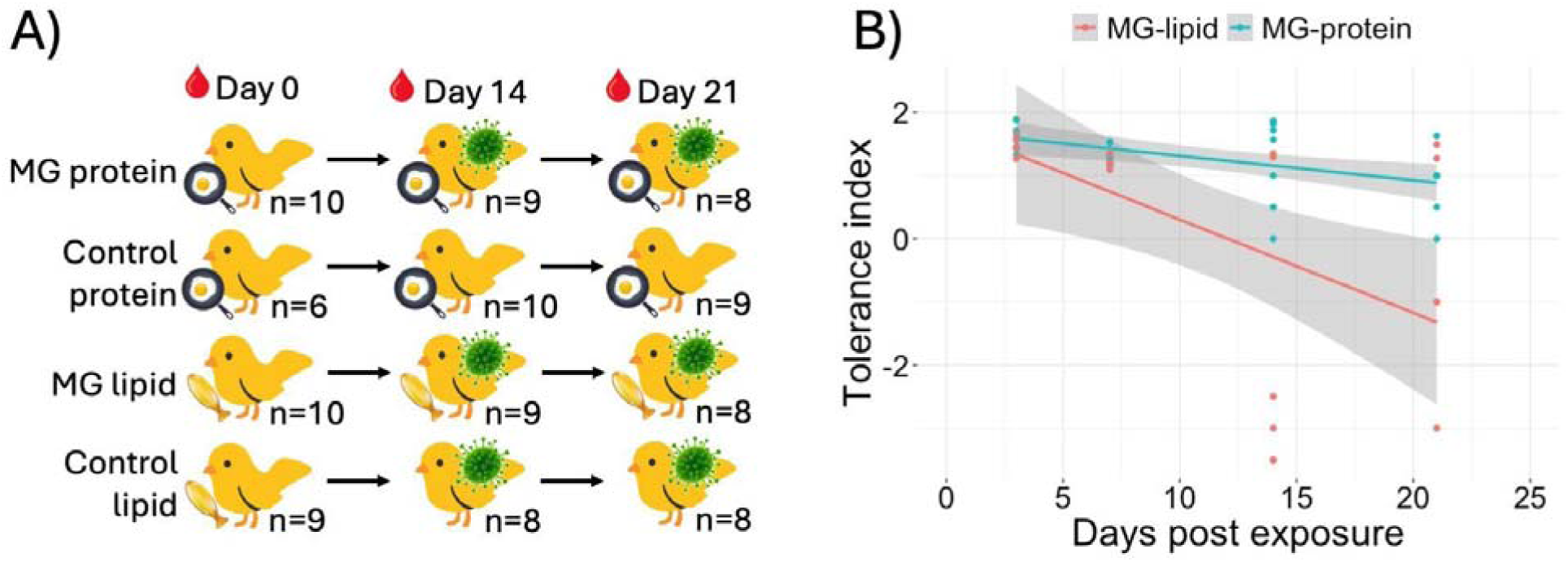
A) Sampling design and sample sizes of domestic canary (*Serinus canaria domestica*) whole blood samples collected during the experiment prior to (day 0) and after *Mycoplasma gallisepticum* (MG)- or sham-exposure (days 14 & 21). Throughout the experiment canaries were fed a diet consisting of a 24 g isocaloric food bar that was high in either protein (20:80 lipid to protein ratio) or lipid content (80:20 lipid to protein ratio). B) Differential effect of high lipid (red points and line) and high protein (blue points and line) diets on canary tolerance to MG infection over time. High protein diet birds were more tolerant of MG infection than birds on the high lipid diet (main effect of diet: = 13.20, *df* = 1, *p* < 0.001; Table S1). Tolerance index is measured as 1-(total eye score/log10(MG load)) where higher values indicate greater tolerance. Points are raw data and gray shading represents associated 95% confidence bands.

## Results and Discussion

### Study design

To examine the main and interactive effects of dietary macronutrients and MG exposure on the transcriptional response of canaries before exposure and during the recovery period, we used edgeR to detect significant differential expression of genes (FDR<0.05) in response to MG, diet, the interaction between diet and infection, and genes correlated to changes in measured phenotypes. A total of 11,124 genes were expressed (cpm > 0.5, mean count > 10) and included in differential expression analysis. One library failed, resulting in a total of 106 sequenced samples (MG-protein: *n* = 27; MG-lipid: *n* = 27; control-protein: *n* = 25; control-lipid: *n* = 25; Figure 1A). Gene Set Enrichment Analysis of KEGG and GO terms for estimated log_2_ fold changes (hereafter logFC) were also conducted for all the comparisons mentioned above.

### Diet drives transcriptional readiness for infection

Changes in gene expression suggest that the macronutrient composition of the diet is integral to the infection response and recovery pattern. These patterns in gene expression also matched diet-driven differences in tolerance. Specifically, protein diet and MG infection exhibit similar effects on gene expression patterns, while lipid diet and MG infection exhibit opposite effects. A total of 497 genes (212 upregulated genes in protein-fed birds vs. 285 upregulated genes in lipid-fed birds, ∼4.6% of genes modeled) among uninfected samples were identified as significantly differentially expressed (FDR<0.05) due to diet (Figures 2A, 2C & S1 & Table S2). Immune response genes were significantly enriched according to diet (GO:0006955, enrichScore=0.70, p.adj=0.02). Immune related ontologies and KEGG pathways similar to those upregulated following MG-exposure are more highly expressed in protein-fed birds (Figure S2). In contrast, lipid-fed birds exhibit a significant functional enrichment for higher expression of the

**Figure 2.**
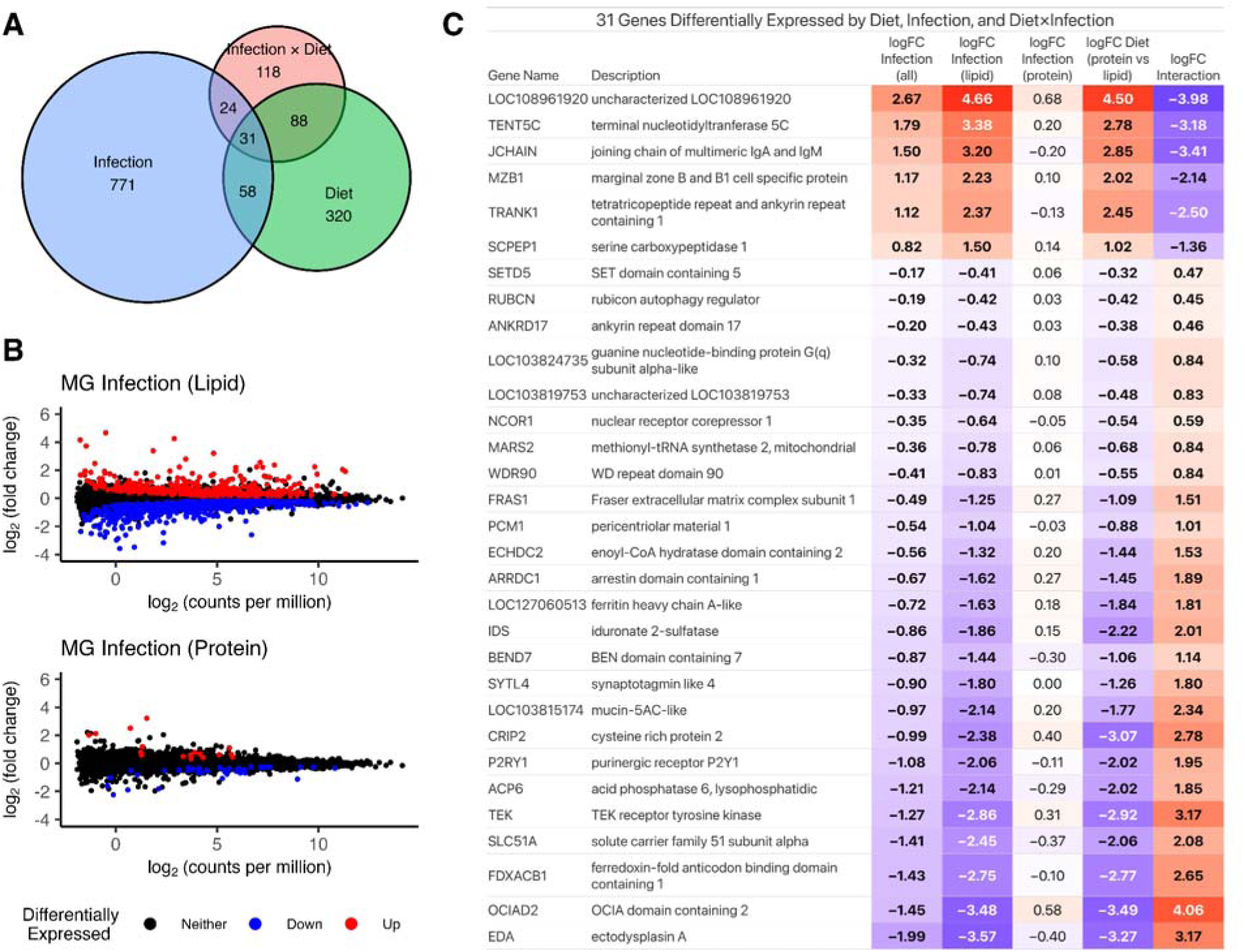
Changes in whole-blood gene expression in response to diet, infection, and their interaction. (A) Total number of genes differentially expressed (FDR < 0.05) in infection, diet and their interaction (see Methods). (B) Density plot of estimated log_2_ fold change (logFC) values for genes differentially expressed following infection in any diet group. Blue and red correspond to logFC estimates for the effect of infection considering only protein-fed and lipid-fed birds, respectively. (C) Table of genes differentially expressed in all three comparisons, bold for FDR<0.05.

Ribosome KEGG pathway (scan03010, enrichScore=-0.47, p.adj<0.04) and extracellular matrix (ECM) receptor interactions (scan04512, enrichScore=-0.66, p.adj<0.02), both of which are downregulated in response to MG infection. This is especially interesting because previous studies suggest early responses to MG are associated with rapidly evolved tolerance in wild house finch populations (31). However, no previous studies have assessed transcriptome differences between tolerant and non-tolerant hosts in the absence of MG, which may be contributing to host capacity to rapidly mitigate damage from MG infection. Here, in the absence of MG, we find that diet-driven differences in functional enrichments suggest transcriptional readiness for infection in protein-fed birds, hence their greater tolerance for infection, and potential immunosuppression in lipid-fed birds.

### Diet-driven transcriptional readiness for infection leads to greater tolerance

After exposure to MG, lipid-fed birds exhibited a more pronounced expression response than protein-fed birds. Thus, MG-induced changes in gene expression following infection are largely driven by the expression responses of lipid-fed diet birds while protein-fed diet birds exhibit comparatively little change in gene expression following infection, connecting diet to changes in transcriptional response to infection (Figure 2B, 2C & S1). This is evident by the notable differences in the distribution of logFC estimates, specifically, there is a bimodal distribution for MG-exposed lipid-fed birds and a zero-centered unimodal distribution for MG-exposed protein-fed birds (Figure 2B & S1). Further, estimated gene expression changes in lipid-fed birds after MG exposure strongly correlate (R = 0.78) with estimated expression changes between diet groups (Figure 3A). In contrast, the gene expression changes following infection for protein-fed birds show a weak negative correlation with expression changes between diet treatment groups (R = −0.45; Figure S3). Indeed, the diet driven patterns of expression and their relationships to tolerance we find in this study are strikingly similar to those found when comparing expression patterns of wild house finch populations that are naive to MG to those that have evolved tolerance to MG (33).

**Figure 3.**
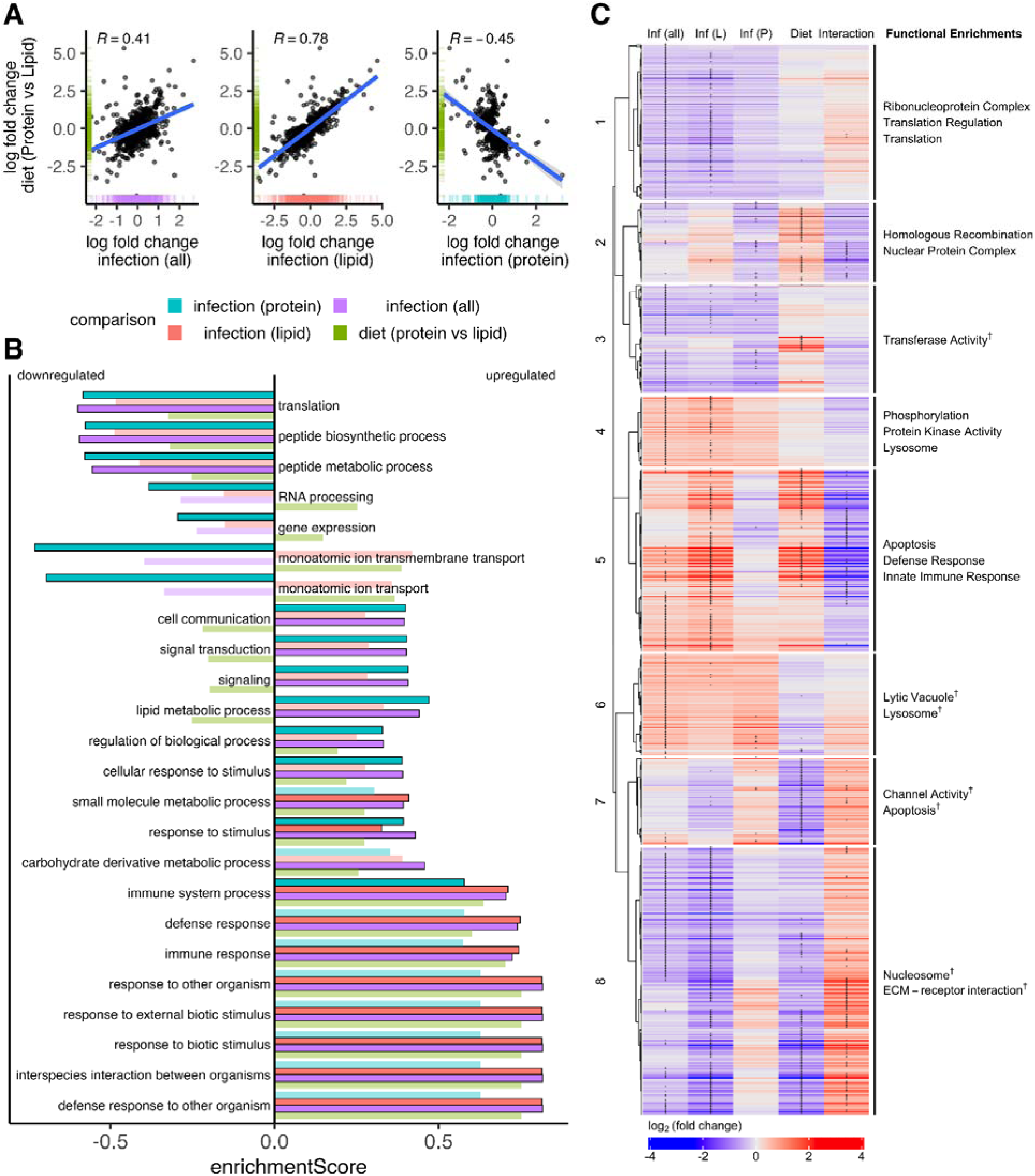
(A) Scatter plots show correlation between estimates for infection by diet and difference by diet among uninfected birds. (B) GO terms enriched for genes changing expression in response to infection. Color corresponds to diet group (red is across all samples, green and blue consider only lipid-fed and protein-fed birds respectively). All terms were significantly enriched in at least one group. Transparent bars correspond to enrichments that did not attain adjusted p-val < 0.05. Colors are shared between panels A and B. (C) Heat map of estimated log_2_ fold change (logFC) by contrast for genes differentially expressed by diet, infection, or the interaction between diet and infection, clustered by logFC similarity. An asterisk is included on genes significant in a given comparison. For infection, red corresponds to increased expression in infection and blue decreased expression. For Diet, red corresponds to higher expression in protein diet and blue higher expression in lipid diet. For interaction, red corresponds to larger relative expression following infection for protein-fed birds than for lipid-fed birds. Contrast wise genes differentially expressed (FDR<0.05) are denoted with x in heatmap. Largest functional enrichments by cluster are shown. The top enrichment terms with p-value < 0.05 for each gene cluster are listed. Ontology terms with FDR > 0.05 denoted with ^†^.

To further elucidate the interaction between diet and infection, differential expression analyses were conducted that compare the difference in the effect of MG on gene expression in protein-fed birds compared to the effect of MG on lipid-fed birds. A total of 261 genes (136 more highly expressed genes in MG-protein birds vs. 125 more highly expressed genes in MG-lipid birds) were identified as significantly differentially expressed (FDR < 0.05) according to diet after MG exposure (Figures 2A, 2C & S1 & Table S2). Immune response (GO:0006955, enrichScore = −0.60, p.adj = 0.13) was the GO term most enriched by interaction (Figure 3B). Increased expression of immune-related genes following infection is observed in lipid-diet relative to protein-fed birds, this is consistent with the enrichments of hierarchical cluster 5 of DE genes (Figure 3C). Surprisingly, infection results in relatively higher mitotic activity of protein-fed birds compared to lipid-fed birds (enrichScore = 0.70, p.adj = 0.27; Figure S2A).

Reduced cell cycle related gene expression in whole blood samples, alongside a relatively smaller increased immune-specific gene expression (GO:0006955, enrichScore = −0.60, p.adj = 0.27), suggests a lipid diet reduces mitotic activity during infection. This is especially striking considering cell-cycle related gene expression is higher in uninfected lipid-fed birds compared to uninfected protein-fed birds. Given MG infection can inhibit cell cycle progression, these results may be due to host immune manipulation by the pathogen (42). KEGG pathway enrichments of the interaction between diet and infection include the ECM-receptor interaction (scan04512), which is downregulated during infection in lipid-fed birds compared to protein-fed birds (enrichScore = 0.64, p.adj = 0.047; Figure S2B). The expression of ECM-receptor genes is significantly less repressed by infection in protein-fed birds compared to lipid-fed birds. Higher expression of ECM receptor related genes is relevant due to the importance of ECM receptors for the cytoadherence and invasion of MG into chicken erythrocytes (43). In contrast, the RIG-I-like receptor signaling pathway is enriched (scan04622, enrichScore = −0.62, p.adj = 0.041) for genes more highly expressed in lipid bird response to infection compared to protein bird response. This enrichment is consistent with the known role of RIG-I receptors in detection of bacterial infections and immune response (44). These functional enrichments for the interaction between diet and MG infection are consistent with known molecular functions. Considering other genes differentially expressed in the interaction between infection and diet reveal a pattern: lipid-fed birds experience enrichment of immune related genes upregulated following infection and downregulation of translational gene expression. These results suggest that diet-based phenotypic differences observed in response to infection occur alongside changes in the transcriptomic expression patterns between diet groups.

### MG infection-induced responses in gene expression are similar to wild finches

Infection served as the largest driver of differential expression. A total of 875 genes (328 up-regulated and 547 down-regulated, ∼7.8% of genes modeled; Figure 2A & Table S2) were identified as significantly differentially expressed (FDR<0.05) when comparing all samples from birds exposed to MG with all unexposed samples. Similar to previous studies on house finches infected with MG (32, 45), most (11 of the 18 terms) of the biological processes enriched for upregulation were immune-specific, including genes related to inflammation, while the 4 terms enriched for downregulation relate to translation *(GO:0006412)* (Figure 3B & Table S3;, 32, 45). Interestingly, the percent of differentially expressed genes in whole blood during the recovery period (∼7.8%) was greater than what other studies have reported from spleen or eye-associated lymphoid tissue during peak infection or during the recovery period (32, 33, 45).

Little is known about the whole-blood gene expression in response to diet or infection in any avian host of MG and this relatively high percent of differentially expressed genes in whole blood is evidence for the utility of blood as a non-destructive and easy to collect tissue sample. Non-destructive sampling is especially useful for understanding host responses over the course of infection or for field sampling wildlife of conservation concern. Further, previous reports of MG invasion of red blood cells occurs in a wide range of hosts (46) and the suggested active role that avian nucleated erythrocytes can play in avian host immunity (47, 48) make the use of whole blood samples for understanding immune responses especially useful.

### Immune and metabolic molecular mechanisms underlie diet-driven tolerance

Genes exhibiting the largest difference between diets in response to infection suggest mechanisms driving divergent immune-related infection responses. Our results provide further evidence that a high lipid-content diet is associated with reduced expression of essential immune-related genes prior to infection (Figure 3A). For example, expression of the Joining Chain Of Multimeric IgA and IgM (JCHAIN) is downregulated by consumption of a high fat diet in mice, resulting in suppressed immunity (49). Our results recapitulate this pattern in many immune genes including JCHAIN, Ig lambda chain C region (LOC103818060), lymphocyte antigen, MYD88 innate immune signal transduction adaptor. Notably, expression of genes that are upregulated in response to infection are already more highly expressed in protein-fed birds compared to lipid-fed birds prior to infection, suggesting repressed immune gene expression prior to infection may be a mechanism for prolonged inflammation in lipid-fed birds. For example, JCHAIN expression prior to infection is 7-fold higher in protein-fed birds compared to lipid-fed birds. In protein-fed birds, no change in JCHAIN expression is observed following infection, while in lipid-fed birds, an additional 9-fold increase JCHAIN expression occurs (Figure 2C). Notably, genes encoding major antibody components dominate genes differentially expressed across interactions. Many of the common immune-related genes were not differentially expressed (interleukins, NF-kB, cytokines), consistent with previous findings that expression of many immune-related genes return to near-baseline 14 days after exposure to MG (34).

Divergent expression of metabolic genes occur in response to MG infection, suggesting a complex interplay between metabolic and immune programming in response to MG infection (Figure 3B). Diet-specific changes in expression of several metabolism-related genes observed following infection may be due to changes in erythrocyte expression or changes in other cell types. Repressed expression of *insulin receptor related receptor (INSRR)* and upregulated *insulin induced gene 2 (INSIG2)* expression was only seen in lipid-fed birds following infection (Figure S4). Persistent dysregulation of INSRR and INSIG2 suggest ongoing metabolic stress. This reasoning is further supported by expression of the *leptin receptor (LEPR)* gene (a type I cytokine receptor), which exhibited a nearly 8-fold divergence in expression patterns following infection between diet groups with downregulation in lipid-fed birds and upregulation in protein-fed birds (FDR=0.006; Table S2). Our results stand in contrast to previous research suggesting that LEPR is unimportant in the avian immune response (50). Differences in metabolic programming between diet groups is not ubiquitous, as both *very low density lipoprotein receptor (VLDLR)* and *lipopolysaccharide induced TNF factor (LITAF)* were upregulated in all birds in response to infection, regardless of diet, suggesting a conserved infection response pattern between diets. Despite some similarities in changes in gene expression, these results suggest that differences in metabolic programming result in increased inflammation for lipid-fed birds, driving observed differences in phenotype.

### Direct assessment of the relationship between tolerance and gene expression

Harnessing the relationships between pathology (conjunctival swelling) phenotype and pathogen load measurements with gene expression partially recapitulates the relationship between MG infection and gene expression while identifying an additional set of genes not previously identified. A total of 130 genes were differentially expressed (FDR<0.05) according to pathogen load (Figure S5, Table S5). Gene set enrichment analysis identified significant repression of translation-related genes as pathogen load increases (GO:0006412, enrichScore = − 0.69, p.adj < 0.001; Figure S6). Increased expression of translational genes at lower levels of pathogen load further suggests MG infection drives translational repression. Adherens junction (scan04520, enrichScore = 0.47, p.adj = 0.02) and lysosome (scan04142, enrichScore = 0.41, p.adj = 0.02) genes are enriched for genes with higher expression with higher pathogen load. Adherens junctions and lysosomes play important immune-related roles in pathogen invasion and clearance, respectively. These results suggest that immune activity in whole-blood samples is observed alongside infection intensity.

The largest overlap in gene lists were those differentially expressed in infection and those that correlate with pathology (i.e., eye score). Meaning, that tolerance (as measured by eye score) and not infection intensity (as measured by pathogen load) drive the infection-induced changes in gene expression (Figure 4A). Ordinal logistic regression was used to identify genes whose expression significantly predicts eye score. A total of 814 genes were identified as significant (FDR<0.05) predictors of conjunctival swelling, with 378 positively correlating and 436 negatively correlating with eye score (Figure S5 & Table S6). Analysis of phenotypic relationships with gene expression indicate that ribosomal gene expression is not only repressed following infection, but the magnitude of repression increases with pathology for many ribosomal genes (GO:0003735, 11/61, p.adj < 0.001; Figure 4B). Mitochondrially localized genes were enriched among genes with expression negatively correlated with eye score (GO: 0005739, 10/69, p.adj < 0.01), but not with infection. Conversely, genes with expression positively correlated with eye score are functionally enriched for ubiquitin-dependent protein catabolism (GO:0006511, 9/38, p.adj < 0.001), a pathway essential for immune ubiquitination activity and regulated cell death. Phenotypic analysis of expression data further supports ribosomal and immune related gene expression exhibit divergent patterns between diet and MG infection, while also suggesting new mechanisms important to the magnitude of response to infection.

**Figure 4.**
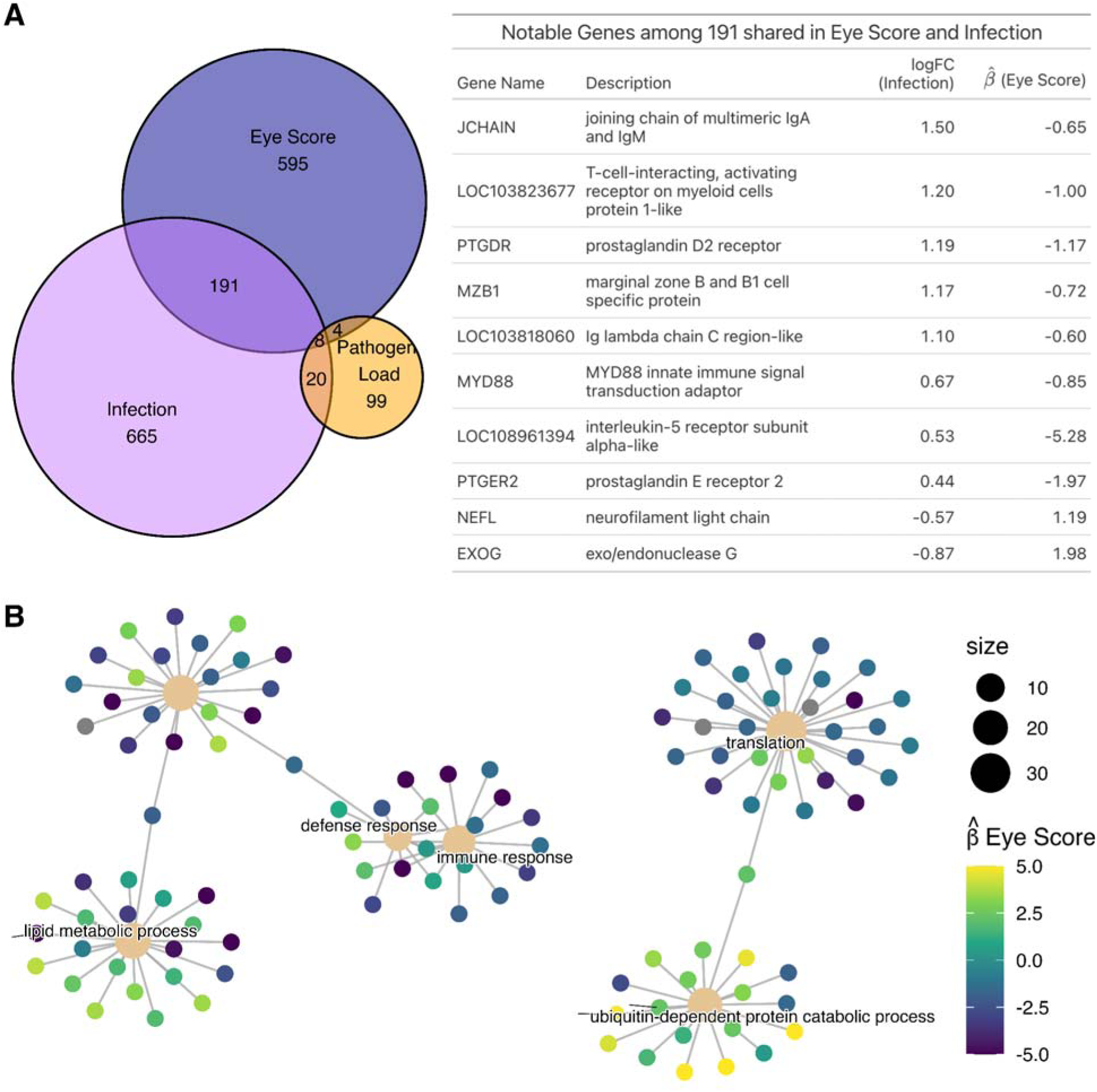
(A) Notable genes that are differentially expressed as both a function of eye score and infection. (B) Venn diagram of genes differentially expressed as a function of eye score, infection, and log_10_ pathogen load. (C) Network plot of top 6 functional enrichments for genes differentially expressed by eye score, with scaled estimated gene beta coefficients plotted for eye score effect.

## Conclusions

We used transcriptomics to understand the currently unexplored effects of diet on gene expression in canary whole blood prior to MG exposure and during the recovery period. As predicted, we found that during the recovery period, many immune and inflammation process genes were upregulated in less tolerant lipid-fed birds compared to protein-fed birds.

Additionally, expression of energy related genes ranging from translation to metabolic pathways were found to be dysregulated as a function of diet and infection. Notably, functional enrichments for genes changing in response to infection are opposite of those for diet, in which genes upregulated in response to infection are higher prior to infection in protein-fed birds, and MG downregulated genes are higher in lipid-fed birds prior to infection. These transcriptome differences between tolerant protein diet and non-tolerant lipid-fed birds in the absence of MG may be contributing to host capacity to rapidly mitigate damage from MG infection (31).

Interestingly, the diet-driven differences in infection response and molecular immune function are strikingly similar to expression patterns seen in wild house finch populations that are naive to MG when compared to those that have evolved tolerance over decades to MG (33). Meaning, diet can cause similar changes to molecular immune mechanisms as those observed under strong pathogen-driven evolutionary selection observed in wild ecosystems. This is especially interesting given the important role of anthropogenic food sources as drivers of transmission in this system and in other wildlife disease systems (1, 7, 8, 25). Considering the substantial effects of diet on immunity, these results provide important insights into the potential for diet to have contributed to the rapid changes in MG tolerance observed in wild bird populations. Finally, these results underscore the importance of considering the role of ecological drivers of host heterogeneity, such as diet, prior to infection and during the recovery process.

## Materials and Methods

### Study species and animal husbandry

MG can infect numerous songbirds species, though it is best studied in house finches (51). We used domestic canaries (*Serinus canaria domestica*) in this experiment as a model for the MG system because canaries are well suited for laboratory studies and continue to exhibit typical life history events in captivity. Canaries and house finches also do not differ in their pathogen loads, production of pathogen-specific serum antibodies, pathology, or recovery time after exposure to identical doses of MG (Hawley et al., 2011).

Further, there is an existing relatively well annotated canary genome (GCF_022539315.1). Canaries were housed individually in cages and provided with either a high protein (*n* = 20) or high lipid (*n* = 22) diet for 17 days prior to inoculation and throughout the experiment. Diets consisted of a 24 g isocaloric food bar that was high in protein (20:80 lipid to protein ratio) or lipid content (80:20 lipid to protein ratio). Both diets contained varying proportions of egg whites, egg yolks, hulled millet, cod liver oil, and were congealed together with agar. On day 0, birds were inoculated in the palpebral conjunctiva of both eyes with either 0.025 mL of MG suspended in Frey’s media (5.00×10^7^ CCU/mL; VA1994; E. Tulman, University of Connecticut) or with a sham of Frey’s media alone. Exposed (lipid: *n* = 13; protein *n* = 10) and sham-exposed individuals (lipid: *n* = 9; protein *n* = 10) were housed on separate racks separated by an opaque room divider to prevent exposure to the pathogen or disease-related social cues (52). For further details regarding husbandry and experiment design see Perrine et al (37).

### Host responses

Disease pathology in canaries was assessed using conjunctival inflammation or “eye score” (0-3 scale per eye, summed to get “total eye score”; modified from Sydenstricker *et al.* 2006). Eye scores were recorded on days 0, 1, 3, 5, 7, then twice a week until day 35. Blood samples (max of 70 µl) and eye swabs were collected prior to exposure and on days 7, 14, and 21. During swabbing, a sterile swab was twirled along the conjunctiva of each eye for 5 sec and stored at −20°C. MG DNA was extracted using Qiagen DNeasy Blood and Tissue protocol (Qiagen, Inc., Valencia, CA) and quantified to determine pathogen load using qPCR methods based on Adelman *et al.* (31) with gBlock® plasmid-based standards (Integrated DNA Technologies, Skokie, IL). All birds were confirmed to be uninfected before the start of the experiment and sham-exposed birds never developed symptoms or tested positive for MG.

### RNA extraction for sequencing

RNAlater was added to blood samples immediately upon collection. Samples were stored at 20°C for 24h then at −80°C until extraction. Canary RNA was extracted from whole blood at days 0, 14, and 21 using a Tri-Reagent phase separation followed by a column clean-up (Direct-zol RNA Miniprep;, 53). RNA was quantified using a Quibit RNA High Sensitivity Assay kit with a Quibit 4 Fluorometer (Thermofisher Scientific, Waltham, MA, USA). Samples with appropriate concentration (7 ng/µl; *N* = 107) were sent to Texas A&M AgriLife Research: Genomics & Bioinformatics Services for randomized QC, library construction, and sequencing. Direction, strand specific RNA libraries were prepared using the PerkinElmer “NEXTflex™ Rapid Directional RNA-Seq Kit 2.0” and the NEXTFLEX® Poly(A) Beads 2.0 kit for Poly-A selection. Libraries were sequenced on two lanes, split equally on a One Novaseq 6000 S2 flowcell (50-bp (2 x 50 bp) paired-end reads). One library failed, resulting in a total of 106 sequenced samples (MG-protein: *n* = 27; MG-lipid: *n* = 27; control-protein: *n* = 25; control-lipid: *n* = 25).

### Tolerance phenotype analysis

To test for differences in infection tolerance (pathology per unit pathogen) over time between diet groups exposed to MG we conducted a generalized linear mixed effects model followed by an ANOVA that included main and interactive effects of diet treatment and time as well as a random intercept for bird identity (Figure 1B; R version 4.1.0 in R Studio; glmmTMB & car packages;, 54–56). Tolerance at individual time points was measured as the ratio between pathology (total eye score) and MG load (log10 transformed) on days 3, 7, 14, and 21 post exposure.

### Bioinformatic analysis

Raw fastq files (GSE273665) were analyzed using the *nf-core* rnaseq workflow version 3.10.1 (57) on the Arkansas High Performance Computing Center computing cluster. The pipeline was executed with Nextflow v22.10.3 (58) utilizing the --star_rsem aligner and --salmon psuedo-aligner.

Reads were screened with fastQC v0.11.9 (59), mapping quality was determined with qualimap v2.2.2 (60), at which point one sample was removed from subsequent analysis due to sequencing failure, with 0.23 million reads passing quality control and 0.02 million mapping to the reference genome. Biotype read proportions were determined via featureCounts of subread software v2.0.1 (61) and read statistics were summarized with RSeQC v3.0.1 (62). Read statistics were compiled into a report with multiQC v1.13 (63), Read trimming was done with Cutadapt v3.4 (64) using default parameters. Trimmed reads were aligned to the serCan2020 refseq gtf reference genome annotation (GCF_022539315.1) with STAR 2.7.10a (65) and counted with RSEM 1.3.1 (66). Detailed mapping statistics are available in the supplemental files (multiqc_report.html). Hemoglobin (Hgb) genes, which comprise a large proportion of RNA reads in whole blood samples, were bioinformatically depleted by removing reads that mapped to genes annotated as Hgb subunits, including LOC103824465, LOC103817747, LOC103824466, LOC103817748, LOC103817749, LOC127059914, and LOC127059976. Exploratory data analysis of count matrices in R was followed by analysis in edgeR (67) using default parameters. Genes with a mean count per million less than 10 were excluded from further analysis via the *filterByExpr* function in edgeR, retaining 10914 genes. Via the function *glmQLFtest(),* the following contrasts were tested:

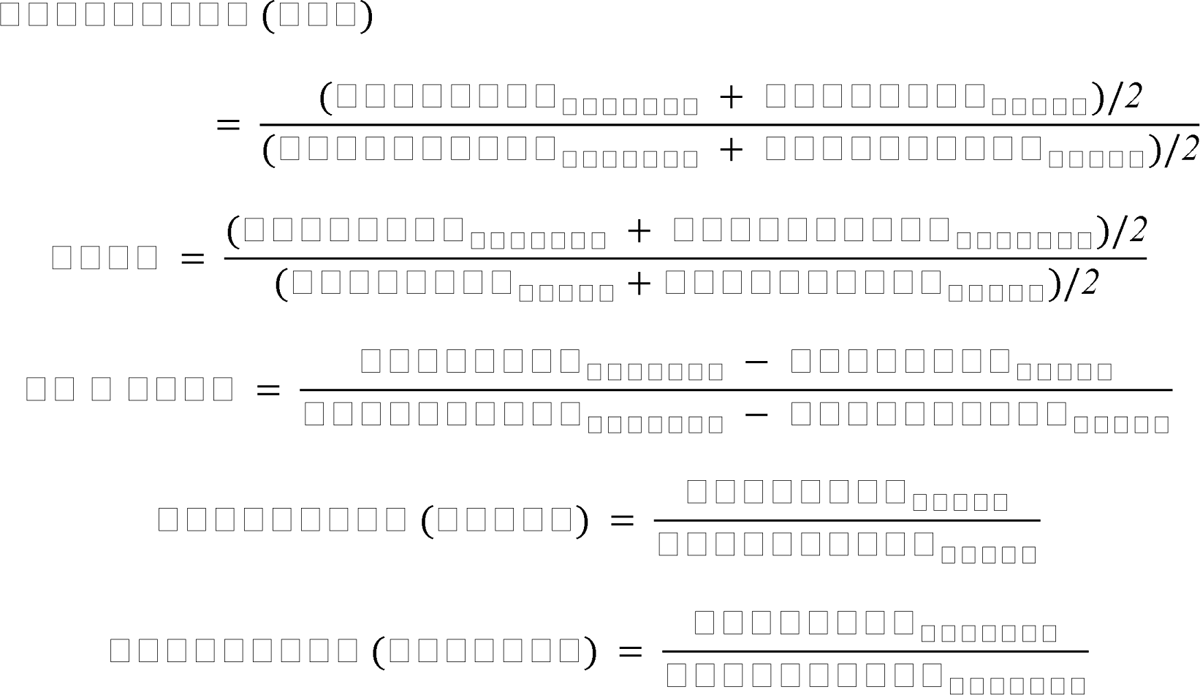

Testing for genes significant in response to phenotypic responses was conducted in edgeR, using each log of pathogen load as a numeric covariate in a regression model, including the effects of diet. Ordinal logistic regression was utilized to estimate effects of each gene on eye score measurements. A model was fit for each gene as EyeScore ∼ logLoad + Diet + {gene_i} using the R package VGAM v1.1-9 (68). Estimated effects of this predictor were used for subsequent analysis, genes were analyzed as described in Pirhaji et al. (69). All p-values noted as p.adjust or FDR have been FDR corrected with the Benjamini & Hochberg method to control false discovery rate. Functional enrichment of differentially expressed genes was conducted using gseGO and gseKEGG in the clusterProfiler package (70). Gene annotations were pulled for “scan” using the AnnotationHub package in. Figures were generated using packages ggplot2, ComplexHeatmap, and ComplexUpset (71–73). Heat map includes all genes identified as significantly differentially expressed in at least one comparison. The number of clusters was determined via the elbow method, implemented in the fviz_nbclust function of the R package factoextra (74). All code used for differential expression analysis and functional enrichment are available in supplemental code.

## Supporting information

Supplemental Results

## Acknowledgments

We thank the Texas A&M AgriLife Research: Genomics & Bioinformatics Services for providing randomized QC, library construction, and sequencing services. Funds were provided by grants to S.E.D. from the National Science Foundation (1941861) and Arkansas Biosciences Institute (Arkansas Settlement Proceeds Act of 2000). Canary experimental methods were approved by the University of Arkansas International Animal Care and Use Committee. This work was supported by resources at the Arkansas High Performance Computing Center, which is funded through multiple National Science Foundation grants and the Arkansas Economic Development Commission.

