## Supplemental Results for "Diet driven differences in host tolerance are linked to shifts in global gene expression in a common avian host-pathogen system"

**Supplemental Figures**

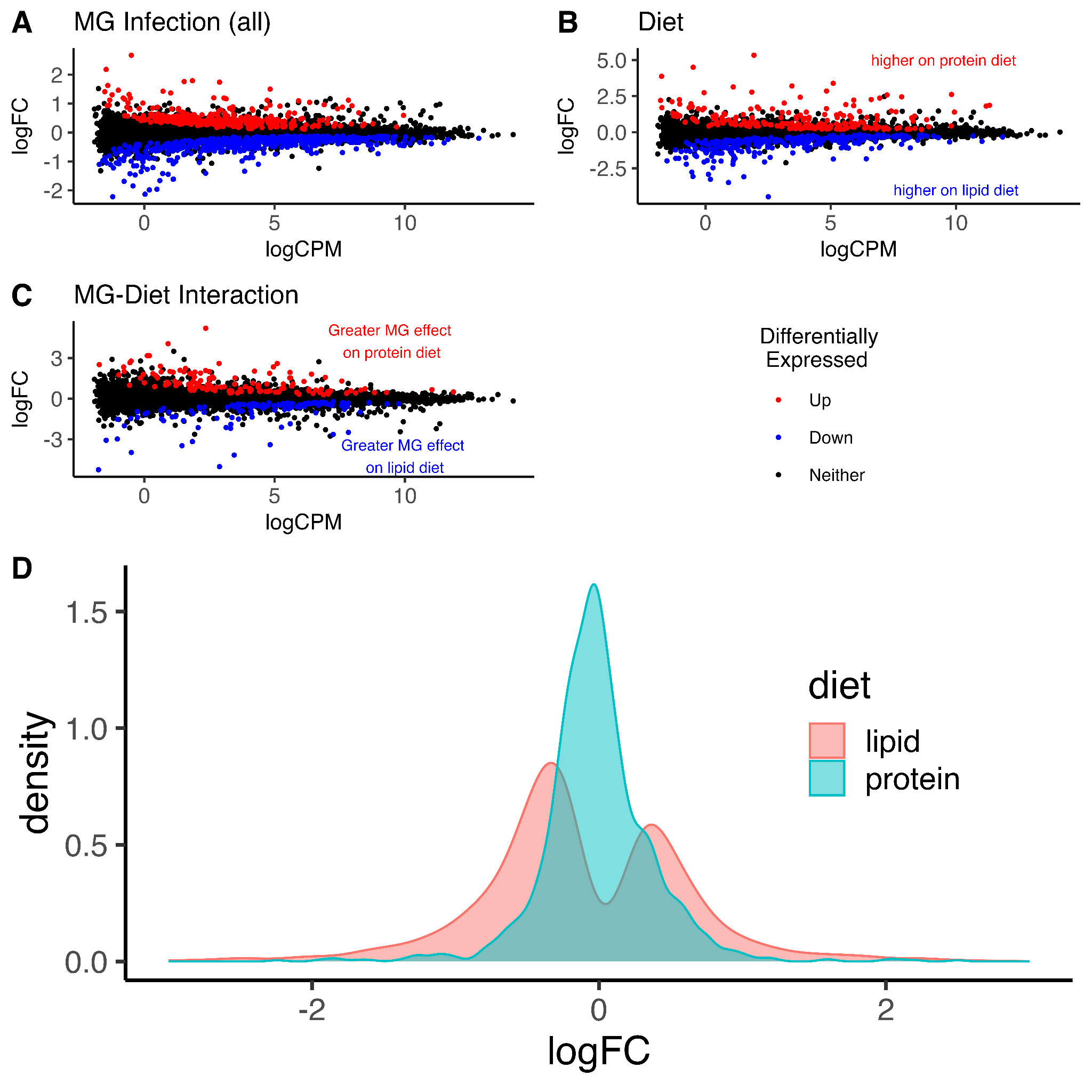

Figure S1: MA plots illustrating differential expression across five contrasts. (A) Effect of MG infection on gene expression. (B) Effect of MG infection on expression in the high lipid diet group. (C) Effect of MG infection on expression in the high protein diet group. (D) The effect of diet (high protein vs. high lipid) on gene expression. (E) Interaction between the effect of MG infection and diet. Each plot displays the log counts per million (logCPM) against the log_2_ fold change (logFC) in expression. Red and blue points denote genes significantly (adjusted p-value < 0.05) upregulated and downregulated respectively, in response to infection unless otherwise noted.

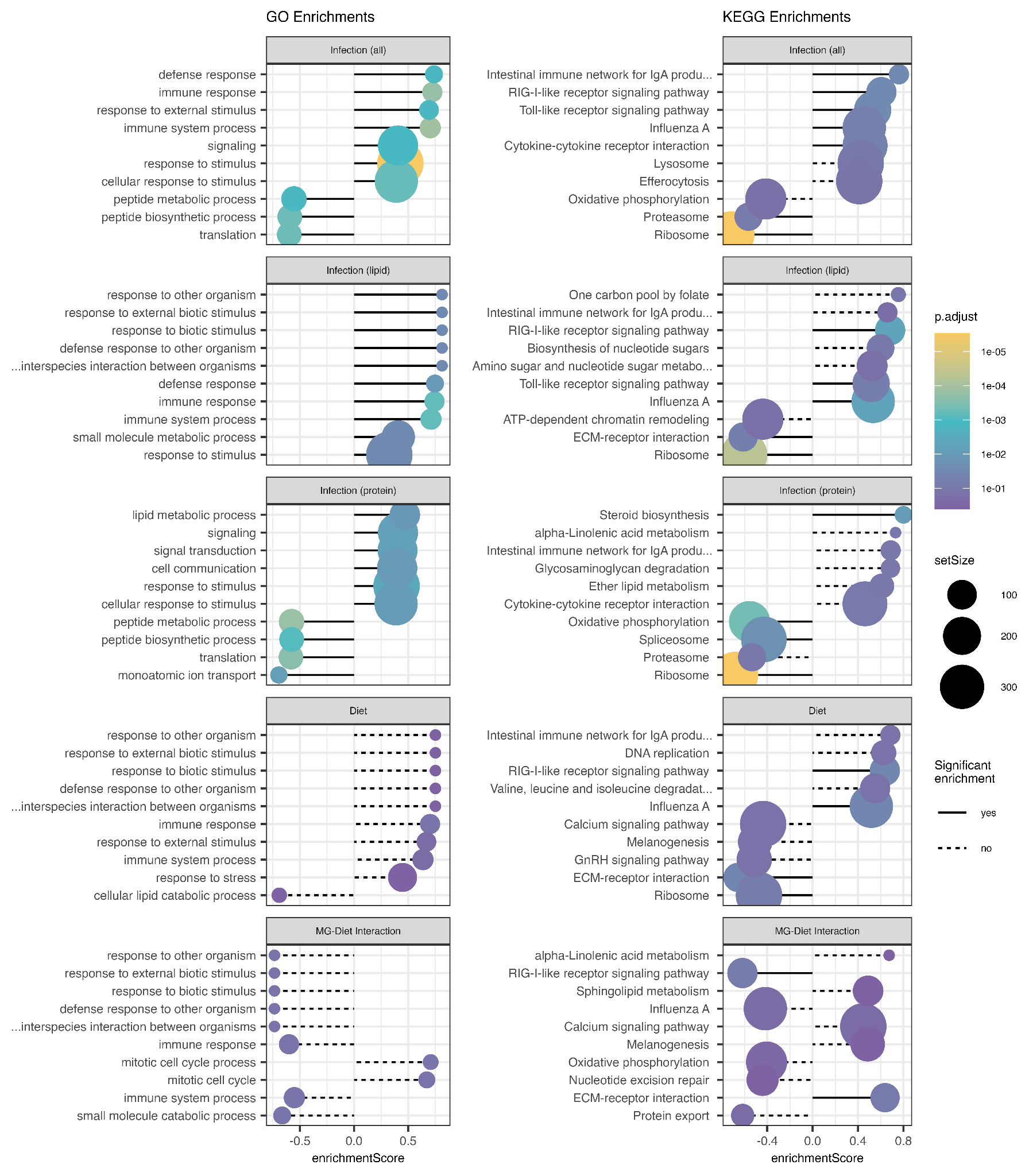

Figure S2: Functional gene sets enriched in gene expression for each contrast. (A) Top 10 Gene Ontology (GO) terms identified in each contrast (B) Top 10 Kyoto Encyclopedia of Genes and Genomes (KEGG) terms identified in each contrast. Each plot displays the log counts per million (logCPM) against the log_2_ fold change (logFC) in expression. Solid lines denote terms significantly (adjusted p-value < 0.05) enriched. Circle size corresponds to the number of genes in that gene set. Circle colors denote the -log10(adjusted p-value). The set enrichmentScore can range from [-1,1], indicating direction of regulation.

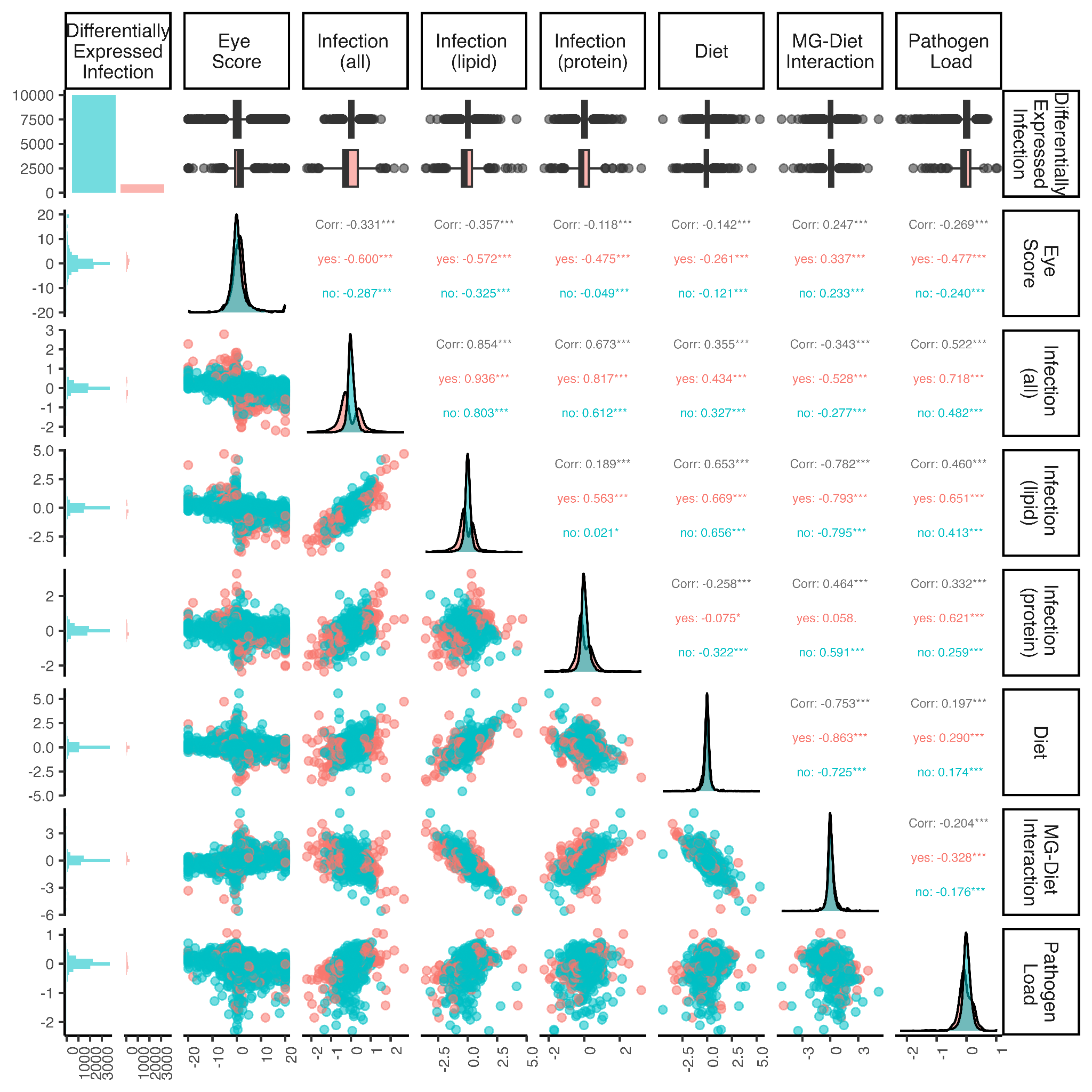

Figure S3: Correlation plot comparing gene-wise expression changes by contrast. Scatter plots in the lower left quadrant show the logFC estimate for each gene. Eye score values denote effects of the gene on eye score logit model. Salmon color data are genes identified as differentially expressed by infection. The upper right quadrant the Pearson correlation coefficient between column and row comparisons, with black text noting value for all cells while salmon and teal colors subset according to significance of the term in the infection contrast. Asterisks denote significance of the correlation, “***” = p <0.001, “**” = p < 0.01, “*”= p < 0.05. Diagonal plots show density of expression change estimates for the corresponding column.

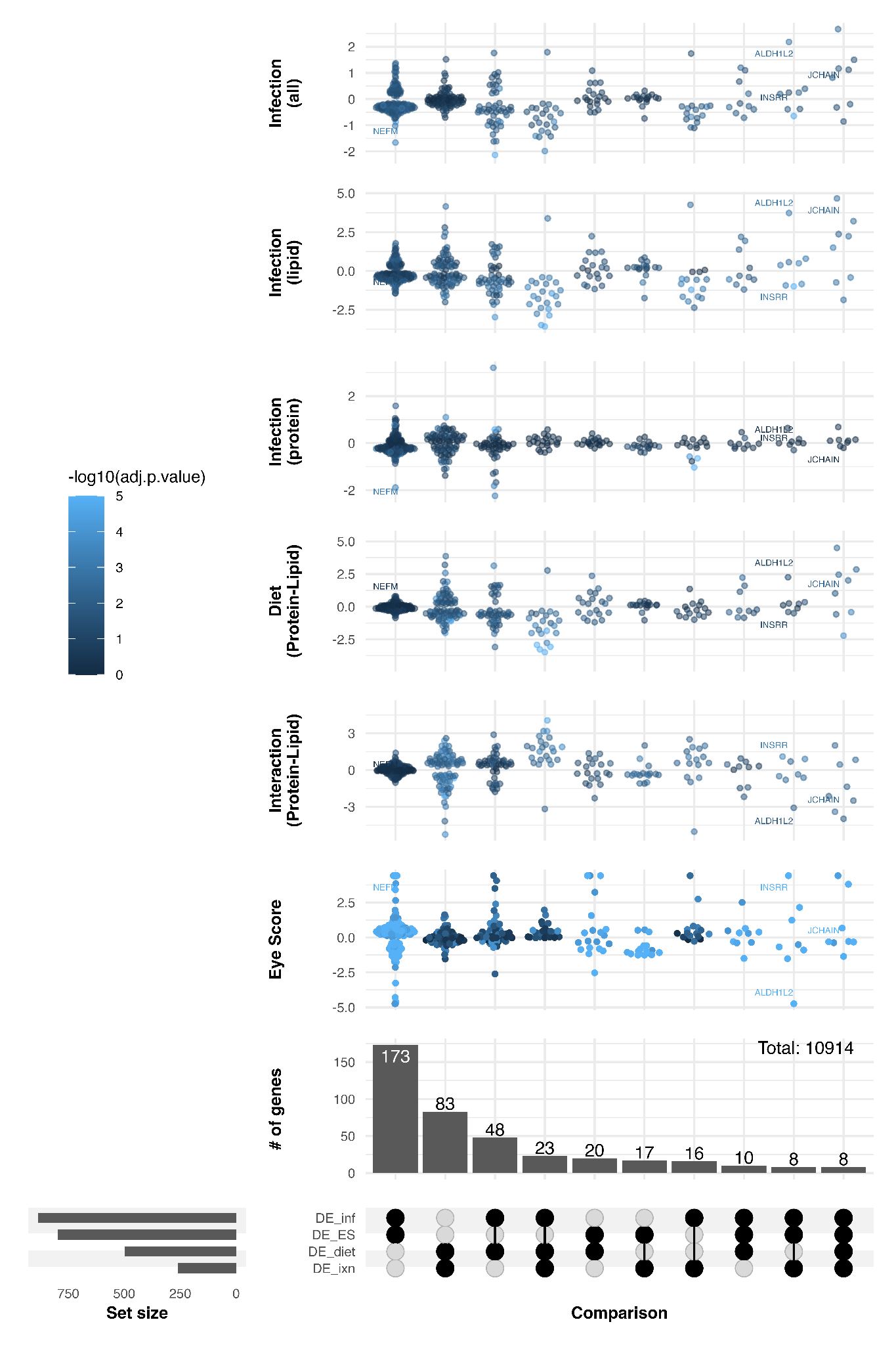

Figure S4: Overlapping differentially expressed genes between each comparison. The number of genes differentially expressed in each comparison are represented by horizontal bars on the left side, while sets of overlap of these genes are tallied on the histogram above comparisons. Beeswarm plots show the distribution of expression changes of the respective set’s genes with one row per comparison.

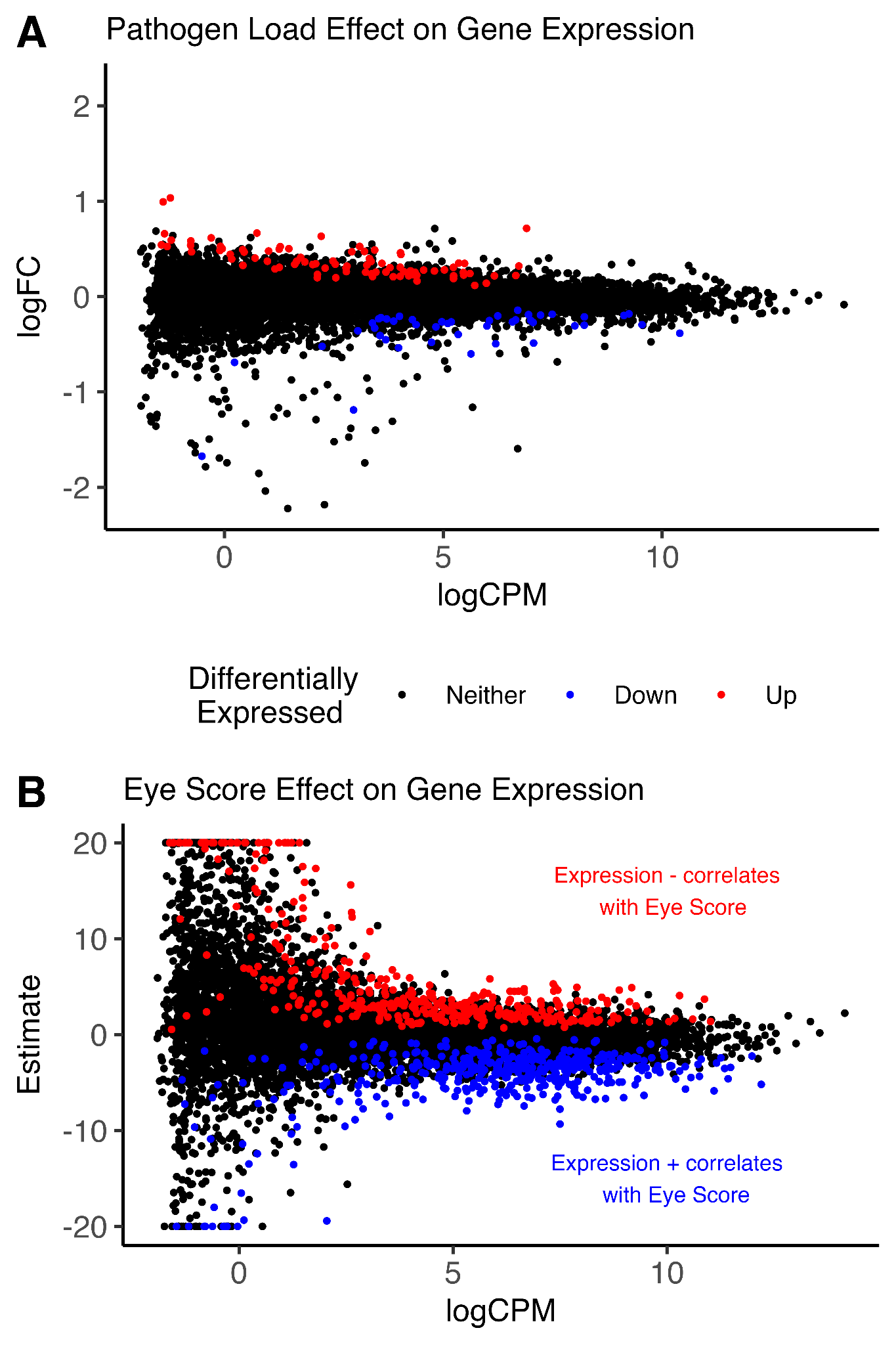

Figure S5: Genes differentially expressed as a function of phenotypic changes, measured in pathogen load or eye score rating. (A) The effect of an increase in the log_10_ pathogen load by one unit detected. Red points correspond to genes whose expression is higher with increasing pathogen load, while blue points are significantly more lowly expressed with increasing pathogen load. (B) The ordinal logistic regression estimates of genes effect on the probability of larger eye scores. Red corresponds to genes whose expression exhibits a significant negative correlation with eye score, while blue gene expression positively correlates with eye score.

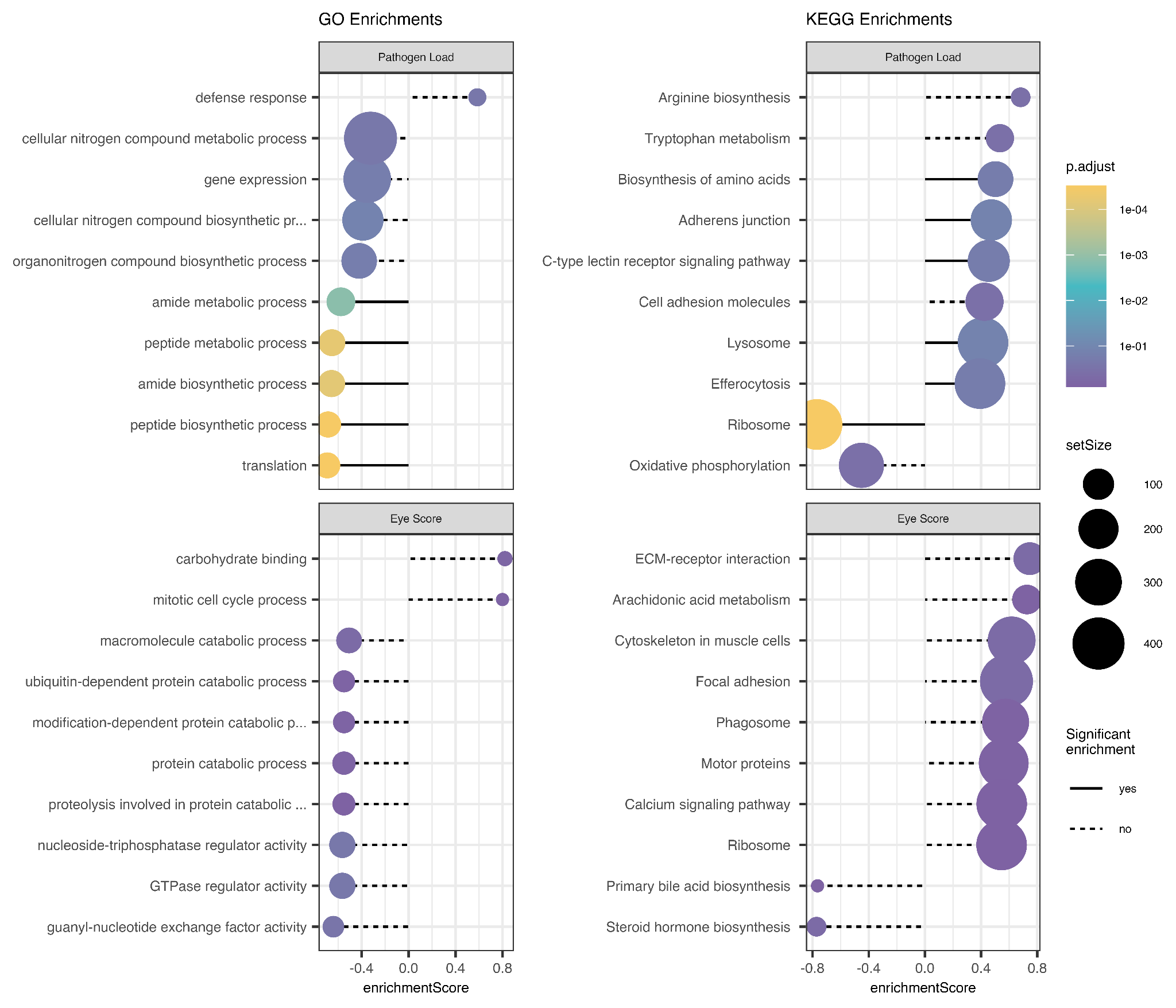

Figure S6: Functional gene sets enriched in gene expression for pathogen load and eye score. (A) Top 10 Gene Ontology (GO) terms identified. (B) Top 10 Kyoto Encyclopedia of Genes and Genomes (KEGG) terms identified. Each plot displays the log counts per million (logCPM) against the log2 fold change (logFC) in expression. Solid lines denote terms significantly (adjusted p-value < 0.05) enriched. Circle size corresponds to the number of genes in that gene set. Circle colors denote the -log10(adjusted p-value). No terms were found to be significant in GSEA for eye score.

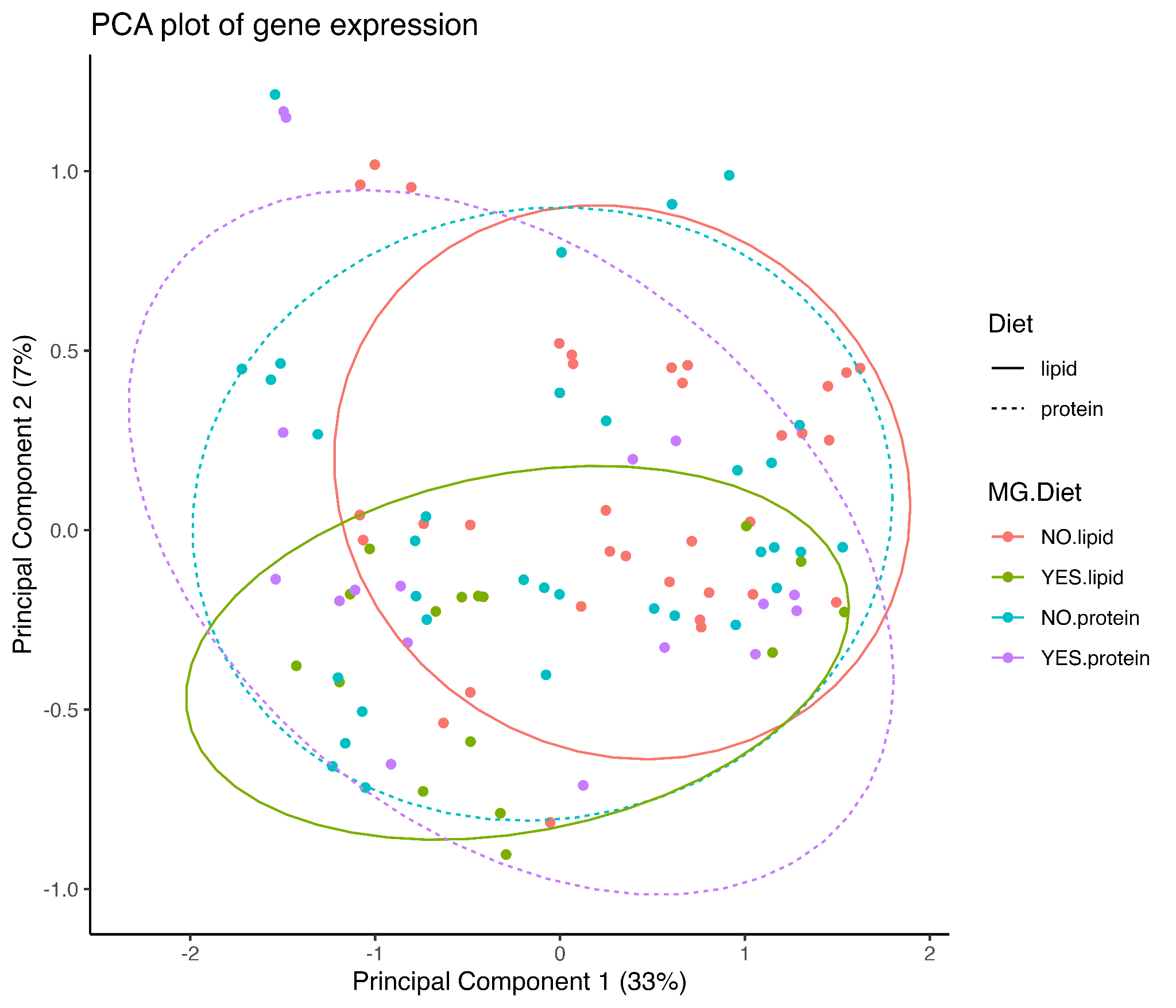

Figure S7: Principal component 1 and 2 scatter plot of the 300 most variable genes. Each point denotes an individual sample. Color denotes MG exposure status and diet treatment group. Ellipses denoted 90% CI for centroid for each group. Solid lines are for lipid-fed groups and dashed lines correspond to protein-fed samples.

**Supplemental Tables**

| Table S1 \| Results from the generalized linear mixed-effects model examining the effect of diet (protein or lipid), days since *Mycoplasma gallisepticum* (MG) exposure, and their interaction on tolerance 1-(total eye score/log10(MG load)) in canaries (*Serinus canaria domestica*) and ANOVA. | | | | |
| --- | --- | --- | --- | --- |
| Generalized linear mixed-effects model | | | | |
|  | Coefficient | SE | z | p value |
| Intercept | 1.780 | 0.434 | 4.098 | <0.0001 |
| DietProtein | -0.080 | 0.584 | -0.137 | 0.891 |
| Day | -0.148 | 0.034 | -4.328 | <0.0001 |
| DietProtein:Day | 0.109 | 0.045 | 2.399 | 0.017 |
| ANOVA | | | | |
|  | Chi squared | | df | p value |
| Diet | 13.196 | | 1 | 0.0003 |
| Day | 14.642 | | 1 | 0.0001 |
| Diet:Day | 5.754 | | 1 | 0.016 |

Table S2: Top 20 genes with the largest magnitude log_2_ fold change in expression for each comparison. MG-Diet interaction genes are genes most differentially expressed in response to infection between diet treatment groups. The MG infection table lists genes with the largest significant magnitude of differential expression between infected and uninfected samples. In diet, positive values correspond to genes more highly expressed in birds on the high protein diet.

| MG-Diet Interaction | | | | |
| --- | --- | --- | --- | --- |
| Top DE Genes | Gene Name | logFC | FDR | description |
| 103824949 | BMP1 | -5.26 | 0.00286 | bone morphogenetic protein 1 |
| 108962424 |  | -5.03 | 0.0125 | uncharacterized LOC108962424 |
| 103821265 |  | -4.17 | 0.0403 | serine/threonine-protein kinase PAK 3-like |
| 108961920 |  | -3.98 | 0.0481 | uncharacterized LOC108961920 |
| 103814621 | LYPD6B | -3.48 | 0.0365 | LY6/PLAUR domain containing 6B |
| 103823426 | JCHAIN | -3.41 | 0.0215 | joining chain of multimeric IgA and IgM |
| 103827129 | NA | -3.18 | 0.037 | NA |
| 103813189 | ALDH1L2 | -3.08 | 0.0453 | aldehyde dehydrogenase 1 family member L2 |
| 127061056 |  | -2.98 | 0.0093 | uncharacterized protein |
| 103815742 | MX1 | -2.65 | 0.0218 | interferon-induced GTP-binding protein Mx-like |
| 103815370 |  | 2.4 | 0.00208 | deoxyribodipyrimidine photo-lyase-like |
| 103818234 | MMP11 | 2.51 | 0.0453 | matrix metallopeptidase 11 |
| 127060989 |  | 2.6 | 0.0436 | uncharacterized LOC127060989 |
| 103818333 | TMEM132B | 2.6 | 0.00677 | transmembrane protein 132B |
| 103823029 | FDXACB1 | 2.65 | 0.0369 | ferredoxin-fold anticodon binding domain containing 1 |
| 103824448 | CRIP2 | 2.78 | 0.0131 | cysteine rich protein 2 |
| 103821834 | TEK | 3.17 | 0.00239 | TEK receptor tyrosine kinase |
| 103816427 | EDA | 3.17 | 0.00676 | ectodysplasin A |
| 103825108 | OCIAD2 | 4.06 | 0.00058 | OCIA domain containing 2 |
| 103826059 | TMEM150C | 5.2 | 0.00562 | transmembrane protein 150C |

| MG Infection | | | | |
| --- | --- | --- | --- | --- |
| Top DE Genes | Gene Name | logFC | FDR | description |
| 103820505 | TTC38 | -2.22 | 0.000438 | tetratricopeptide repeat domain 38 |
| 103815985 | CGNL1 | -2.13 | 2.01e-05 | cingulin like 1 |
| 103816427 | EDA | -1.99 | 0.000377 | ectodysplasin A |
| 103822998 | POU2F3 | -1.96 | 0.00158 | POU class 2 homeobox 3 |
| 103812707 |  | -1.85 | 0.00765 | sodium channel protein type 5 subunit alpha-like |
| 103815634 |  | -1.69 | 0.0162 | phospholipase A and acyltransferase 1-like |
| 103822758 | NEFM | -1.66 | 0.000377 | neurofilament medium chain |
| 103818123 | FAM222A | -1.62 | 0.0146 | family with sequence similarity 222 member A |
| 103826188 | KIT | -1.61 | 0.00407 | KIT proto-oncogene, receptor tyrosine kinase |
| 127059203 |  | -1.47 | 0.0284 | feather beta keratin-like |
| 115484236 | SMIM33 | 1.41 | 0.0208 | small integral membrane protein 33 |
| 103814454 | KLHL38 | 1.43 | 0.00461 | kelch like family member 38 |
| 103826749 | ARHGAP28 | 1.46 | 0.00776 | Rho GTPase activating protein 28 |
| 103823426 | JCHAIN | 1.5 | 0.0249 | joining chain of multimeric IgA and IgM |
| 127061143 |  | 1.63 | 0.00881 | cathepsin S-like |
| 108962424 |  | 1.74 | 0.0491 | uncharacterized LOC108962424 |
| 103817978 | IQCD | 1.76 | 0.0139 | IQ motif containing D |
| 103827129 | NA | 1.79 | 0.00903 | NA |
| 103813189 | ALDH1L2 | 2.18 | 0.00181 | aldehyde dehydrogenase 1 family member L2 |
| 108961920 |  | 2.67 | 0.00407 | uncharacterized LOC108961920 |

| Diet (Protein-Lipid) | | | | |
| --- | --- | --- | --- | --- |
| Top DE Genes | Gene Name | logFC | FDR | description |
| 103822244 |  | -4.47 | 6.74e-05 | mas-related G-protein coupled receptor member H-like |
| 103825108 | OCIAD2 | -3.49 | 5.88e-06 | OCIA domain containing 2 |
| 103816427 | EDA | -3.27 | 5.88e-06 | ectodysplasin A |
| 103817978 | IQCD | -3.09 | 0.00535 | IQ motif containing D |
| 103824448 | CRIP2 | -3.07 | 7.88e-06 | cysteine rich protein 2 |
| 103821834 | TEK | -2.92 | 5.94e-06 | TEK receptor tyrosine kinase |
| 103823029 | FDXACB1 | -2.77 | 0.000409 | ferredoxin-fold anticodon binding domain containing 1 |
| 127061044 |  | -2.47 | 0.00286 | centrosome-associated protein CEP250-like |
| 103812751 | NA | -2.23 | 0.00832 | NA |
| 103816359 | IDS | -2.22 | 0.000111 | iduronate 2-sulfatase |
| 103824282 | OASL | 2.61 | 0.00284 | 2'-5'-oligoadenylate synthase 1-like |
| 103826266 | SLC39A8 | 2.74 | 0.000337 | solute carrier family 39 member 8 |
| 103827129 | NA | 2.78 | 0.00846 | NA |
| 103823426 | JCHAIN | 2.85 | 0.00459 | joining chain of multimeric IgA and IgM |
| 103818123 | FAM222A | 3.13 | 0.000588 | family with sequence similarity 222 member A |
| 103821265 |  | 3.2 | 0.0244 | serine/threonine-protein kinase PAK 3-like |
| 103818339 | RIMBP2 | 3.38 | 0.000253 | RIMS binding protein 2 |
| 103824949 | BMP1 | 3.87 | 0.00293 | bone morphogenetic protein 1 |
| 108961920 |  | 4.5 | 0.00262 | uncharacterized LOC108961920 |
| 103822244 |  | -4.47 | 6.74e-05 | mas-related G-protein coupled receptor member H-like |

Table S3: Gene Ontology terms for each comparison tested, sorted by adjusted p values. Set Size corresponds to the number of genes annotated to the GO term for S. canaria. Enrich Score, taking values from -1 to 1, denotes the relative localization of these terms relative to the entire transcriptome, with a large positive corresponding to enriched for positive fold changes in expression, and large negative enriched among genes with negative log fold change terms.

| **Comparison** | **ID** | **Description** | **Set**  **Size** | **Enrich**  **Score** | **p.adjust** |
| --- | --- | --- | --- | --- | --- |
| Infection (all) | GO:0050896 | response to stimulus | 338 | 0.428 | 2.90e-06 |
|  | GO:0002376 | immune system process | 37 | 0.705 | 1.13e-04 |
|  | GO:0006955 | immune response | 32 | 0.724 | 1.64e-04 |
|  | GO:0006412 | translation | 58 | -0.600 | 5.35e-04 |
|  | GO:0043043 | peptide biosynthetic process | 59 | -0.594 | 5.35e-04 |
|  | GO:0051716 | cellular response to stimulus | 281 | 0.391 | 6.04e-04 |
|  | GO:0023052 | signaling | 231 | 0.407 | 1.19e-03 |
|  | GO:0009605 | response to external stimulus | 30 | 0.694 | 1.34e-03 |
|  | GO:0006518 | peptide metabolic process | 64 | -0.555 | 1.34e-03 |
|  | GO:0006952 | defense response | 23 | 0.739 | 1.36e-03 |
|  | GO:0007165 | signal transduction | 227 | 0.402 | 1.38e-03 |
|  | GO:0007154 | cell communication | 233 | 0.396 | 1.38e-03 |
|  | GO:0043604 | amide biosynthetic process | 66 | -0.531 | 1.71e-03 |
|  | GO:0050789 | regulation of biological process | 427 | 0.331 | 2.41e-03 |
|  | GO:0065007 | biological regulation | 440 | 0.339 | 2.56e-03 |
|  | GO:0009607 | response to biotic stimulus | 12 | 0.817 | 3.03e-03 |
|  | GO:0043207 | response to external biotic stimulus | 12 | 0.817 | 3.03e-03 |
|  | GO:0044419 | biological process involved in interspecies interaction between organisms | 12 | 0.817 | 3.03e-03 |
|  | GO:0051707 | response to other organism | 12 | 0.817 | 3.03e-03 |
|  | GO:0098542 | defense response to other organism | 12 | 0.817 | 3.03e-03 |
|  | GO:0006950 | response to stress | 94 | 0.486 | 3.16e-03 |
|  | GO:0006954 | inflammatory response | 13 | 0.778 | 7.58e-03 |
|  | GO:0050794 | regulation of cellular process | 408 | 0.328 | 8.57e-03 |
|  | GO:0043603 | amide metabolic process | 78 | -0.464 | 1.93e-02 |
|  | GO:0006629 | lipid metabolic process | 108 | 0.441 | 2.34e-02 |
|  | GO:1901135 | carbohydrate derivative metabolic process | 81 | 0.458 | 3.04e-02 |
|  | GO:0044281 | small molecule metabolic process | 141 | 0.393 | 3.04e-02 |
|  | GO:0002682 | regulation of immune system process | 13 | 0.729 | 3.16e-02 |
|  | GO:0048584 | positive regulation of response to stimulus | 22 | 0.620 | 6.41e-02 |
|  | GO:0016042 | lipid catabolic process | 30 | 0.570 | 6.41e-02 |
| Diet (Protein -Lipid) | GO:0006955 | immune response | 32 | 0.703 | 1.30e-01 |
|  | GO:0009605 | response to external stimulus | 30 | 0.669 | 2.19e-01 |
|  | GO:0002376 | immune system process | 37 | 0.637 | 2.19e-01 |
|  | GO:0009607 | response to biotic stimulus | 12 | 0.753 | 4.10e-01 |
|  | GO:0043207 | response to external biotic stimulus | 12 | 0.753 | 4.10e-01 |
|  | GO:0044419 | biological process involved in interspecies interaction between organisms | 12 | 0.753 | 4.10e-01 |
|  | GO:0051707 | response to other organism | 12 | 0.753 | 4.10e-01 |
|  | GO:0098542 | defense response to other organism | 12 | 0.753 | 4.10e-01 |
|  | GO:0044242 | cellular lipid catabolic process | 16 | -0.691 | 4.10e-01 |
|  | GO:0006950 | response to stress | 94 | 0.448 | 4.10e-01 |
|  | GO:0040011 | locomotion | 15 | 0.671 | 6.81e-01 |
|  | GO:0006952 | defense response | 23 | 0.601 | 6.81e-01 |
|  | GO:0006935 | chemotaxis | 14 | 0.665 | 6.81e-01 |
|  | GO:0042330 | taxis | 14 | 0.665 | 6.81e-01 |
|  | GO:0042981 | regulation of apoptotic process | 26 | 0.557 | 6.81e-01 |
|  | GO:0043067 | regulation of programmed cell death | 26 | 0.557 | 6.81e-01 |
|  | GO:0006259 | DNA metabolic process | 70 | 0.427 | 6.81e-01 |
|  | GO:0006897 | endocytosis | 16 | -0.625 | 8.84e-01 |
|  | GO:0098657 | import into cell | 16 | -0.625 | 8.84e-01 |
|  | GO:0080134 | regulation of response to stress | 17 | 0.600 | 8.84e-01 |
|  | GO:0007167 | enzyme-linked receptor protein signaling pathway | 11 | -0.663 | 8.84e-01 |
|  | GO:0043547 | positive regulation of GTPase activity | 10 | 0.674 | 8.84e-01 |
|  | GO:0051345 | positive regulation of hydrolase activity | 12 | 0.632 | 8.84e-01 |
|  | GO:0043087 | regulation of GTPase activity | 12 | 0.630 | 8.84e-01 |
|  | GO:0051336 | regulation of hydrolase activity | 20 | 0.562 | 8.84e-01 |
|  | GO:0043414 | macromolecule methylation | 17 | 0.567 | 8.84e-01 |
|  | GO:0006633 | fatty acid biosynthetic process | 10 | -0.634 | 8.84e-01 |
|  | GO:0072330 | monocarboxylic acid biosynthetic process | 10 | -0.634 | 8.84e-01 |
|  | GO:0006511 | ubiquitin-dependent protein catabolic process | 37 | 0.470 | 8.84e-01 |
|  | GO:0019941 | modification-dependent protein catabolic process | 37 | 0.470 | 8.84e-01 |
| MG-Diet Interaction | GO:0044282 | small molecule catabolic process | 23 | -0.664 | 1.28e-01 |
|  | GO:0006955 | immune response | 32 | -0.602 | 1.28e-01 |
|  | GO:0009605 | response to external stimulus | 30 | -0.587 | 1.28e-01 |
|  | GO:0009607 | response to biotic stimulus | 12 | -0.736 | 1.28e-01 |
|  | GO:0043207 | response to external biotic stimulus | 12 | -0.736 | 1.28e-01 |
|  | GO:0044419 | biological process involved in interspecies interaction between organisms | 12 | -0.736 | 1.28e-01 |
|  | GO:0051707 | response to other organism | 12 | -0.736 | 1.28e-01 |
|  | GO:0098542 | defense response to other organism | 12 | -0.736 | 1.28e-01 |
|  | GO:0006952 | defense response | 23 | -0.632 | 1.28e-01 |
|  | GO:0002376 | immune system process | 37 | -0.552 | 1.28e-01 |
|  | GO:0051603 | proteolysis involved in protein catabolic process | 41 | -0.539 | 1.28e-01 |
|  | GO:1903047 | mitotic cell cycle process | 17 | 0.709 | 1.28e-01 |
|  | GO:0006511 | ubiquitin-dependent protein catabolic process | 37 | -0.531 | 1.28e-01 |
|  | GO:0019941 | modification-dependent protein catabolic process | 37 | -0.531 | 1.28e-01 |
|  | GO:0000278 | mitotic cell cycle | 20 | 0.673 | 1.28e-01 |
|  | GO:0016579 | protein deubiquitination | 20 | -0.638 | 1.32e-01 |
|  | GO:0009260 | ribonucleotide biosynthetic process | 15 | -0.695 | 1.43e-01 |
|  | GO:0046390 | ribose phosphate biosynthetic process | 15 | -0.695 | 1.43e-01 |
|  | GO:0034220 | monoatomic ion transmembrane transport | 13 | -0.707 | 1.43e-01 |
|  | GO:0030163 | protein catabolic process | 43 | -0.510 | 1.43e-01 |
|  | GO:0007167 | enzyme-linked receptor protein signaling pathway | 11 | 0.751 | 1.43e-01 |
|  | GO:0016054 | organic acid catabolic process | 17 | -0.648 | 1.51e-01 |
|  | GO:0046395 | carboxylic acid catabolic process | 17 | -0.648 | 1.51e-01 |
|  | GO:0070646 | protein modification by small protein removal | 21 | -0.619 | 1.52e-01 |
|  | GO:0009259 | ribonucleotide metabolic process | 19 | -0.607 | 1.55e-01 |
|  | GO:0019693 | ribose phosphate metabolic process | 19 | -0.607 | 1.55e-01 |
|  | GO:1901565 | organonitrogen compound catabolic process | 61 | -0.447 | 1.67e-01 |
|  | GO:0022402 | cell cycle process | 34 | 0.543 | 1.67e-01 |
|  | GO:1901575 | organic substance catabolic process | 115 | -0.376 | 1.67e-01 |
|  | GO:0006811 | monoatomic ion transport | 21 | -0.597 | 1.97e-01 |
| Infection (protein) | GO:0006518 | peptide metabolic process | 64 | -0.578 | 1.52e-04 |
|  | GO:0006412 | translation | 58 | -0.583 | 2.48e-04 |
|  | GO:0043043 | peptide biosynthetic process | 59 | -0.577 | 8.70e-04 |
|  | GO:0050896 | response to stimulus | 338 | 0.393 | 3.46e-03 |
|  | GO:0023052 | signaling | 231 | 0.407 | 6.82e-03 |
|  | GO:0007165 | signal transduction | 227 | 0.403 | 6.82e-03 |
|  | GO:0006811 | monoatomic ion transport | 21 | -0.695 | 7.24e-03 |
|  | GO:0051716 | cellular response to stimulus | 281 | 0.389 | 8.85e-03 |
|  | GO:0007154 | cell communication | 233 | 0.399 | 9.46e-03 |
|  | GO:0006629 | lipid metabolic process | 108 | 0.470 | 1.15e-02 |
|  | GO:0043603 | amide metabolic process | 78 | -0.454 | 1.33e-02 |
|  | GO:0043604 | amide biosynthetic process | 66 | -0.479 | 1.65e-02 |
|  | GO:0010467 | gene expression | 317 | -0.295 | 2.16e-02 |
|  | GO:0006396 | RNA processing | 105 | -0.384 | 3.23e-02 |
|  | GO:0034220 | monoatomic ion transmembrane transport | 13 | -0.730 | 3.76e-02 |
|  | GO:0002376 | immune system process | 37 | 0.578 | 3.76e-02 |
|  | GO:0016042 | lipid catabolic process | 30 | 0.596 | 3.76e-02 |
|  | GO:0000375 | RNA splicing, via transesterification reactions | 32 | -0.541 | 4.15e-02 |
|  | GO:0000377 | RNA splicing, via transesterification reactions with bulged adenosine as nucleophile | 32 | -0.541 | 4.15e-02 |
|  | GO:0000398 | mRNA splicing, via spliceosome | 32 | -0.541 | 4.15e-02 |
|  | GO:0006812 | monoatomic cation transport | 13 | -0.709 | 4.76e-02 |
|  | GO:0030001 | metal ion transport | 13 | -0.709 | 4.76e-02 |
|  | GO:0034470 | ncRNA processing | 43 | -0.485 | 4.76e-02 |
|  | GO:0034660 | ncRNA metabolic process | 55 | -0.452 | 4.76e-02 |
|  | GO:0050789 | regulation of biological process | 427 | 0.329 | 4.76e-02 |
|  | GO:0022613 | ribonucleoprotein complex biogenesis | 42 | -0.491 | 5.04e-02 |
|  | GO:0065007 | biological regulation | 440 | 0.328 | 5.04e-02 |
|  | GO:0006955 | immune response | 32 | 0.574 | 5.08e-02 |
|  | GO:0050794 | regulation of cellular process | 408 | 0.329 | 5.36e-02 |
|  | GO:0006954 | inflammatory response | 13 | 0.694 | 9.47e-02 |
| Infection (lipid) | GO:0006955 | immune response | 32 | 0.744 | 6.52e-04 |
|  | GO:0002376 | immune system process | 37 | 0.711 | 6.52e-04 |
|  | GO:0006952 | defense response | 23 | 0.749 | 1.04e-02 |
|  | GO:0009607 | response to biotic stimulus | 12 | 0.814 | 2.96e-02 |
|  | GO:0043207 | response to external biotic stimulus | 12 | 0.814 | 2.96e-02 |
|  | GO:0044419 | biological process involved in interspecies interaction between organisms | 12 | 0.814 | 2.96e-02 |
|  | GO:0051707 | response to other organism | 12 | 0.814 | 2.96e-02 |
|  | GO:0098542 | defense response to other organism | 12 | 0.814 | 2.96e-02 |
|  | GO:0044281 | small molecule metabolic process | 141 | 0.409 | 2.96e-02 |
|  | GO:0050896 | response to stimulus | 338 | 0.326 | 3.01e-02 |
|  | GO:0009605 | response to external stimulus | 30 | 0.649 | 5.13e-02 |
|  | GO:0044282 | small molecule catabolic process | 23 | 0.656 | 1.08e-01 |
|  | GO:0006954 | inflammatory response | 13 | 0.724 | 1.23e-01 |
|  | GO:0016054 | organic acid catabolic process | 17 | 0.680 | 1.27e-01 |
|  | GO:0046395 | carboxylic acid catabolic process | 17 | 0.680 | 1.27e-01 |
|  | GO:0006950 | response to stress | 94 | 0.417 | 1.30e-01 |
|  | GO:1903047 | mitotic cell cycle process | 17 | -0.667 | 1.48e-01 |
|  | GO:0002682 | regulation of immune system process | 13 | 0.699 | 1.48e-01 |
|  | GO:0048584 | positive regulation of response to stimulus | 22 | 0.611 | 1.48e-01 |
|  | GO:0043043 | peptide biosynthetic process | 59 | -0.487 | 1.48e-01 |
|  | GO:0006412 | translation | 58 | -0.484 | 1.48e-01 |
|  | GO:0006082 | organic acid metabolic process | 77 | 0.432 | 1.48e-01 |
|  | GO:0019752 | carboxylic acid metabolic process | 77 | 0.432 | 1.48e-01 |
|  | GO:0043436 | oxoacid metabolic process | 77 | 0.432 | 1.48e-01 |
|  | GO:0043604 | amide biosynthetic process | 66 | -0.466 | 1.48e-01 |
|  | GO:0019637 | organophosphate metabolic process | 72 | 0.432 | 2.00e-01 |
|  | GO:0090407 | organophosphate biosynthetic process | 46 | 0.493 | 2.38e-01 |
|  | GO:0043087 | regulation of GTPase activity | 12 | 0.684 | 2.38e-01 |
|  | GO:0000278 | mitotic cell cycle | 20 | -0.617 | 2.38e-01 |
|  | GO:0009967 | positive regulation of signal transduction | 12 | 0.682 | 2.38e-01 |
| Pathogen Load | GO:0006412 | translation | 58 | -0.691 | 3.01e-05 |
|  | GO:0043043 | peptide biosynthetic process | 59 | -0.685 | 3.08e-05 |
|  | GO:0006518 | peptide metabolic process | 64 | -0.652 | 6.01e-05 |
|  | GO:0043604 | amide biosynthetic process | 66 | -0.655 | 6.90e-05 |
|  | GO:0043603 | amide metabolic process | 78 | -0.576 | 1.32e-03 |
|  | GO:0044271 | cellular nitrogen compound biosynthetic process | 216 | -0.389 | 1.20e-01 |
|  | GO:0010467 | gene expression | 317 | -0.352 | 1.59e-01 |
|  | GO:1901566 | organonitrogen compound biosynthetic process | 144 | -0.420 | 1.61e-01 |
|  | GO:0034641 | cellular nitrogen compound metabolic process | 418 | -0.324 | 2.33e-01 |
|  | GO:0006952 | defense response | 23 | 0.585 | 2.40e-01 |
|  | GO:0009607 | response to biotic stimulus | 12 | 0.672 | 3.61e-01 |
|  | GO:0043207 | response to external biotic stimulus | 12 | 0.672 | 3.61e-01 |
|  | GO:0044419 | biological process involved in interspecies interaction between organisms | 12 | 0.672 | 3.61e-01 |
|  | GO:0051707 | response to other organism | 12 | 0.672 | 3.61e-01 |
|  | GO:0098542 | defense response to other organism | 12 | 0.672 | 3.61e-01 |
|  | GO:0005975 | carbohydrate metabolic process | 44 | 0.456 | 3.71e-01 |
|  | GO:0009059 | macromolecule biosynthetic process | 361 | -0.311 | 3.80e-01 |
|  | GO:0043547 | positive regulation of GTPase activity | 10 | 0.670 | 3.94e-01 |
|  | GO:0051345 | positive regulation of hydrolase activity | 12 | 0.646 | 4.83e-01 |
|  | GO:0006954 | inflammatory response | 13 | 0.615 | 4.83e-01 |
|  | GO:1901576 | organic substance biosynthetic process | 456 | -0.293 | 4.83e-01 |
|  | GO:0042592 | homeostatic process | 12 | -0.696 | 5.31e-01 |
|  | GO:0048856 | anatomical structure development | 42 | -0.495 | 5.31e-01 |
|  | GO:0007049 | cell cycle | 75 | -0.430 | 5.31e-01 |
|  | GO:0044249 | cellular biosynthetic process | 442 | -0.295 | 5.31e-01 |
|  | GO:0009058 | biosynthetic process | 464 | -0.289 | 5.31e-01 |
|  | GO:0006633 | fatty acid biosynthetic process | 10 | -0.699 | 5.36e-01 |
|  | GO:0072330 | monocarboxylic acid biosynthetic process | 10 | -0.699 | 5.36e-01 |
|  | GO:0008610 | lipid biosynthetic process | 44 | -0.491 | 5.48e-01 |
|  | GO:1901605 | alpha-amino acid metabolic process | 14 | 0.586 | 5.48e-01 |
| Eye Score | GO:0030695 | GTPase regulator activity | 60 | -0.564 | 2.36e-01 |
|  | GO:0060589 | nucleoside-triphosphatase regulator activity | 60 | -0.564 | 2.36e-01 |
|  | GO:0005085 | guanyl-nucleotide exchange factor activity | 35 | -0.640 | 2.78e-01 |
|  | GO:0009057 | macromolecule catabolic process | 57 | -0.506 | 4.74e-01 |
|  | GO:0030246 | carbohydrate binding | 18 | 0.820 | 6.16e-01 |
|  | GO:1903047 | mitotic cell cycle process | 17 | 0.799 | 7.00e-01 |
|  | GO:0030163 | protein catabolic process | 43 | -0.550 | 7.00e-01 |
|  | GO:0006511 | ubiquitin-dependent protein catabolic process | 37 | -0.549 | 7.00e-01 |
|  | GO:0019941 | modification-dependent protein catabolic process | 37 | -0.549 | 7.00e-01 |
|  | GO:0051603 | proteolysis involved in protein catabolic process | 41 | -0.550 | 7.80e-01 |
|  | GO:0022402 | cell cycle process | 34 | 0.689 | 7.80e-01 |
|  | GO:0051301 | cell division | 39 | 0.661 | 8.22e-01 |
|  | GO:0000278 | mitotic cell cycle | 20 | 0.768 | 8.54e-01 |
|  | GO:0005911 | cell-cell junction | 12 | 0.814 | 9.10e-01 |
|  | GO:0010498 | proteasomal protein catabolic process | 14 | -0.656 | 1.00e+00 |
|  | GO:0048285 | organelle fission | 11 | 0.803 | 1.00e+00 |
|  | GO:0020037 | heme binding | 13 | 0.773 | 1.00e+00 |
|  | GO:0043632 | modification-dependent macromolecule catabolic process | 38 | -0.514 | 1.00e+00 |
|  | GO:0070161 | anchoring junction | 21 | 0.715 | 1.00e+00 |
|  | GO:0043161 | proteasome-mediated ubiquitin-dependent protein catabolic process | 11 | -0.664 | 1.00e+00 |
|  | GO:0046903 | secretion | 11 | 0.774 | 1.00e+00 |
|  | GO:0030054 | cell junction | 40 | 0.632 | 1.00e+00 |
|  | GO:0005681 | spliceosomal complex | 21 | -0.556 | 1.00e+00 |
|  | GO:0008186 | ATP-dependent activity, acting on RNA | 14 | -0.608 | 1.00e+00 |
|  | GO:0003724 | RNA helicase activity | 13 | -0.616 | 1.00e+00 |
|  | GO:0046906 | tetrapyrrole binding | 14 | 0.716 | 1.00e+00 |
|  | GO:0007049 | cell cycle | 75 | 0.574 | 1.00e+00 |
|  | GO:0004930 | G protein-coupled receptor activity | 60 | 0.583 | 1.00e+00 |
|  | GO:0099080 | supramolecular complex | 46 | 0.603 | 1.00e+00 |
|  | GO:0016055 | Wnt signaling pathway | 13 | -0.600 | 1.00e+00 |

Table S4: Functional enrichment of genes in each cluster of the heat map of differentially expressed genes in Figure 3C of the main text. The five GO and KEGG terms enrichments with the smallest p-values are listed below, representing enrichments of the genes included in that cluster.

| **Cluster** | **ID** | **Description** | **Gene Ratio** | **Background Ratio** | **p.adjust** |
| --- | --- | --- | --- | --- | --- |
| 1 | GO:0032991 | protein-containing complex | 17/39 | 420/2702 | 0.00144 |
|  | GO:0008135 | translation factor activity, RNA binding | 4/41 | 15/2722 | 0.00255 |
|  | GO:0090079 | translation regulator activity, nucleic acid binding | 4/41 | 15/2722 | 0.00255 |
|  | GO:0045182 | translation regulator activity | 4/41 | 17/2722 | 0.0029 |
|  | GO:1990904 | ribonucleoprotein complex | 6/39 | 84/2702 | 0.0319 |
|  | scan03010 | Ribosome | 12/79 | 109/3881 | 0.000119 |
|  | scan04810 | Regulation of actin cytoskeleton | 7/79 | 145/3881 | 0.871 |
|  | scan00564 | Glycerophospholipid metabolism | 4/79 | 65/3881 | 0.871 |
|  | scan04910 | Insulin signaling pathway | 5/79 | 104/3881 | 0.871 |
|  | scan04530 | Tight junction | 5/79 | 108/3881 | 0.871 |
| 2 | GO:0140513 | nuclear protein-containing complex | 5/19 | 94/2702 | 0.021 |
|  | GO:0031981 | nuclear lumen | 4/19 | 84/2702 | 0.0684 |
|  | GO:0016043 | cellular component organization | 5/12 | 236/2159 | 0.371 |
|  | GO:0031974 | membrane-enclosed lumen | 4/19 | 116/2702 | 0.0879 |
|  | GO:0043233 | organelle lumen | 4/19 | 116/2702 | 0.0879 |
|  | scan04141 | Protein processing in endoplasmic reticulum | 4/28 | 133/3881 | 0.2 |
| 3 | GO:0016740 | transferase activity | 9/20 | 479/2722 | 0.184 |
|  | GO:0043412 | macromolecule modification | 8/19 | 371/2159 | 0.361 |
|  | GO:0016772 | transferase activity, transferring phosphorus-containing groups | 5/20 | 220/2722 | 0.184 |
|  | GO:0004672 | protein kinase activity | 4/20 | 148/2722 | 0.184 |
|  | GO:0016070 | RNA metabolic process | 6/19 | 265/2159 | 0.361 |
|  | scan01240 | Biosynthesis of cofactors | 4/52 | 98/3881 | 0.707 |
| 4 | GO:0004672 | protein kinase activity | 6/17 | 148/2722 | 0.0127 |
|  | GO:0006468 | protein phosphorylation | 6/15 | 150/2159 | 0.0214 |
|  | GO:0016773 | phosphotransferase activity, alcohol group as acceptor | 6/17 | 167/2722 | 0.0127 |
|  | GO:0016301 | kinase activity | 6/17 | 184/2722 | 0.0143 |
|  | GO:0016310 | phosphorylation | 6/15 | 175/2159 | 0.0249 |
|  | scan04142 | Lysosome | 6/39 | 109/3881 | 0.0321 |
| 5 | GO:0045087 | innate immune response | 4/50 | 15/2159 | 0.0402 |
|  | GO:0009607 | response to biotic stimulus | 4/50 | 20/2159 | 0.0402 |
|  | GO:0043207 | response to external biotic stimulus | 4/50 | 20/2159 | 0.0402 |
|  | GO:0044419 | biological process involved in interspecies interaction between organisms | 4/50 | 20/2159 | 0.0402 |
|  | GO:0051707 | response to other organism | 4/50 | 20/2159 | 0.0402 |
|  | scan05164 | Influenza A | 11/86 | 93/3881 | 0.00044 |
|  | scan04210 | Apoptosis | 10/86 | 103/3881 | 0.00354 |
|  | scan04115 | p53 signaling pathway | 5/86 | 55/3881 | 0.222 |
|  | scan03018 | RNA degradation | 4/86 | 62/3881 | 0.566 |
|  | scan04620 | Toll-like receptor signaling pathway | 4/86 | 65/3881 | 0.566 |
| 6 | GO:0006629 | lipid metabolic process | 5/19 | 145/2159 | 0.508 |
|  | GO:0044255 | cellular lipid metabolic process | 4/19 | 96/2159 | 0.508 |
|  | GO:0050896 | response to stimulus | 9/19 | 487/2159 | 0.508 |
|  | GO:0016787 | hydrolase activity | 10/29 | 494/2722 | 0.272 |
|  | GO:0140657 | ATP-dependent activity | 4/29 | 118/2722 | 0.281 |
|  | scan04142 | Lysosome | 6/55 | 109/3881 | 0.22 |
|  | scan04621 | NOD-like receptor signaling pathway | 4/55 | 96/3881 | 0.509 |
|  | scan04010 | MAPK signaling pathway | 4/55 | 165/3881 | 0.618 |
| 7 | GO:0032991 | protein-containing complex | 5/19 | 420/2702 | 0.643 |
|  | GO:0000166 | nucleotide binding | 4/13 | 468/2722 | 0.467 |
|  | GO:1901265 | nucleoside phosphate binding | 4/13 | 468/2722 | 0.467 |
| 8 | GO:0044085 | cellular component biogenesis | 10/48 | 139/2159 | 0.165 |
|  | GO:0000786 | nucleosome | 4/61 | 22/2702 | 0.0876 |
|  | GO:0022607 | cellular component assembly | 8/48 | 114/2159 | 0.356 |
|  | GO:0000785 | chromatin | 4/61 | 37/2702 | 0.135 |
|  | GO:0043228 | non-membrane-bounded organelle | 14/61 | 317/2702 | 0.135 |
|  | scan04512 | ECM-receptor interaction | 5/119 | 37/3881 | 0.352 |
|  | scan04912 | GnRH signaling pathway | 6/119 | 57/3881 | 0.352 |
|  | scan04916 | Melanogenesis | 5/119 | 54/3881 | 0.753 |
|  | scan04310 | Wnt signaling pathway | 6/119 | 96/3881 | 0.898 |
|  | scan04540 | Gap junction | 4/119 | 54/3881 | 0.898 |

Table S5: Genes differentially expressed as a function of pathogen load. The 20 largest magnitude positive and negative genes with FDR < 0.05, ordered by log_2_ fold change.

| **Top DE Genes** | **Gene Name** | **logFC** | **FDR** | **Description** |
| --- | --- | --- | --- | --- |
| 103814123 | VWA2 | -1.67 | 0.0267 | von Willebrand factor A domain containing 2 |
| 103827276 |  | -1.19 | 4.67e-05 | mitochondrial nicotinamide adenine dinucleotide transporter SLC25A51-like |
| 103822983 | PTS | -0.69 | 0.043 | 6-pyruvoyltetrahydropterin synthase |
| 127060226 |  | -0.601 | 0.000863 | uncharacterized LOC127060226 |
| 103820377 | IRAK4 | -0.537 | 0.0106 | interleukin 1 receptor associated kinase 4 |
| 103813962 | BLOC1S2 | -0.522 | 0.00114 | biogenesis of lysosomal organelles complex 1 subunit 2 |
| 103819314 | FAM161A | -0.495 | 0.0283 | FAM161 centrosomal protein A |
| 103812846 | TMEM106B | -0.488 | 0.0134 | transmembrane protein 106B |
| 127060478 |  | -0.48 | 0.0106 | uncharacterized LOC127060478 |
| 103812504 | CEP295 | -0.453 | 0.0448 | centrosomal protein 295 |
| 103820796 | ATAD3A | -0.409 | 0.00393 | ATPase family AAA domain containing 3A |
| 103822559 | TRNAU1AP | -0.397 | 0.0263 | tRNA selenocysteine 1 associated protein 1 |
| 103824732 |  | -0.385 | 0.0106 | ubiquitin-associated protein 2-like |
| 103814543 | HIBCH | -0.359 | 0.0448 | 3-hydroxyisobutyryl-CoA hydrolase |
| 103812513 | FAM76B | -0.331 | 0.0267 | family with sequence similarity 76 member B |
| 103827273 | CEP57 | -0.318 | 0.0317 | centrosomal protein 57 |
| 103822322 | TSTD3 | -0.309 | 0.0397 | thiosulfate sulfurtransferase like domain containing 3 |
| 103815219 | AK4 | -0.307 | 0.0448 | adenylate kinase 4 |
| 103824667 | MTHFD2 | -0.3 | 0.0415 | methylenetetrahydrofolate dehydrogenase (NADP+ dependent) 2, methenyltetrahydrofolate cyclohydrolase |
| 103827423 | GTF2F2 | -0.296 | 0.0356 | general transcription factor IIF subunit 2 |
| 103815614 | SLC12A9 | 0.488 | 0.0356 | solute carrier family 12 member 9 |
| 103819747 | BAIAP2 | 0.491 | 0.0282 | BAR/IMD domain containing adaptor protein 2 |
| 108963641 | ATP6V1B1 | 0.497 | 0.0386 | ATPase H+ transporting V1 subunit B1 |
| 103825600 | SLC39A1 | 0.502 | 0.00126 | solute carrier family 39 member 1 |
| 103822527 | RNF150 | 0.502 | 0.0106 | ring finger protein 150 |
| 115484288 |  | 0.523 | 0.011 | nascent polypeptide-associated complex subunit alpha, muscle-specific form-like |
| 103813692 | CADPS2 | 0.525 | 0.0495 | calcium dependent secretion activator 2 |
| 103822711 | RETSAT | 0.525 | 0.0223 | retinol saturase |
| 108963022 |  | 0.526 | 0.0106 | heat shock protein 30C-like |
| 103817376 | SH3PXD2B | 0.538 | 0.0448 | SH3 and PX domains 2B |
| 127061154 |  | 0.542 | 0.0363 | integrin alpha-5-like |
| 127059608 |  | 0.583 | 0.0146 | uncharacterized LOC127059608 |
| 127059127 |  | 0.591 | 0.0142 | uncharacterized LOC127059127 |
| 103822809 | KCNJ1 | 0.615 | 0.0356 | potassium inwardly rectifying channel subfamily J member 1 |
| 103819001 | ASS1 | 0.632 | 0.0315 | argininosuccinate synthase 1 |
| 108964832 | FCRL6 | 0.659 | 0.0356 | Fc receptor like 6 |
| 103819231 | PAK5 | 0.666 | 0.000935 | p21 (RAC1) activated kinase 5 |
| 103814579 | FN1 | 0.715 | 0.00906 | fibronectin 1 |
| 127061143 |  | 0.993 | 0.002 | cathepsin S-like |
| 103813113 | FZD1 | 1.03 | 4.67e-05 | frizzled class receptor 1 |

Table S6: Genes with the largest estimated effect on eye score phenotype among genes with p.adjust < 0.05. Estimate corresponds to the estimated beta coefficient in the ordinal logistic regression model for that gene. The p value corresponds to the p-value for that estimate. Positive estimate values correspond to higher likelihood of a lower eye score.

| **Top DE Genes** | **Gene Name** | **Estimate** | **Pr(>\|z\|)** | **description** |
| --- | --- | --- | --- | --- |
| 103813189 | ALDH1L2 | -22.9 | 0.000191 | aldehyde dehydrogenase 1 family member L2 |
| 103819791 | ATAD5 | -19.4 | 4.71e-05 | ATPase family AAA domain containing 5 |
| 103813815 | NHLRC2 | -7.95 | 0.000138 | NHL repeat containing 2 |
| 103821685 | ROPN1L | -7.2 | 0.000135 | rhophilin associated tail protein 1 like |
| MARCHF6 | MARCHF6 | -6.57 | 0.000102 | membrane associated ring-CH-type finger 6 |
| 103826516 | GPATCH2L | -6.51 | 0.000195 | G-patch domain containing 2 like |
| 103813947 | MAPK8 | -6.49 | 0.000195 | mitogen-activated protein kinase 8 |
| 103826195 | LONRF1 | -6.24 | 0.000116 | LON peptidase N-terminal domain and ring finger 1 |
| 103826382 |  | -5.92 | 0.000105 | SERTA domain-containing protein 2-like |
| 103826536 | FBXO33 | -5.52 | 3.84e-05 | F-box protein 33 |
| 103815744 | ZBTB21 | -5.51 | 0.00013 | zinc finger and BTB domain containing 21 |
| 103822284 | CNTF | -5.32 | 0.000112 | ciliary neurotrophic factor |
| 103826471 | ATXN3 | -5.17 | 7.34e-05 | ataxin 3 |
| 103812931 | NR1D2 | -5.02 | 3.21e-05 | nuclear receptor subfamily 1 group D member 2 |
| 103813515 | HBP1 | -4.35 | 0.000128 | HMG-box transcription factor 1 |
| 103818075 | YPEL1 | -3.63 | 7.31e-05 | yippee like 1 |
| 103818267 | SERINC1 | -3.4 | 1.63e-05 | serine incorporator 1 |
| 103826521 | FOS | -2.82 | 8.12e-05 | Fos proto-oncogene, AP-1 TF subunit |
| 103818289 | CRYBB3 | -2.45 | 3.22e-06 | crystallin beta B3 |
| 103823426 | JCHAIN | -0.648 | 0.000122 | joining chain of multimeric IgA and IgM |
| 103815780 | MID1IP1 | 1.9 | 0.000603 | MID1 interacting protein 1 |
| 103827408 | ITM2B | 2.71 | 7.72e-05 | integral membrane protein 2B |
| 103818732 | PSMA7 | 3.01 | 0.000468 | proteasome 20S subunit alpha 7 |
| 103826500 | SLIRP | 3.29 | 0.000518 | SRA stem-loop interacting RNA binding protein |
| 103815819 | ASB9 | 3.41 | 0.000443 | ankyrin repeat and SOCS box containing 9 |
| 103821870 | TIMM13 | 3.43 | 0.00067 | translocase of inner mitochondrial membrane 13 |
| 103815935 | RPS17 | 3.54 | 0.000303 | ribosomal protein S17 |
| 103822338 | KCTD20 | 3.65 | 0.000281 | potassium channel tetramerization domain containing 20 |
| 103820311 | PIGL | 4.04 | 0.000263 | phosphatidylinositol glycan anchor biosynthesis class L |
| 103822424 |  | 4.07 | 0.000272 | glutathione S-transferase 2 |
| 103814116 | PRDX3 | 4.3 | 0.000269 | peroxiredoxin 3 |
| 103816717 | KATNB1 | 4.51 | 0.000434 | katanin regulatory subunit B1 |
| 103823572 | RHCE | 4.64 | 0.000185 | Rh blood group CcEe antigens |
| 103823724 | HKDC1 | 5.77 | 0.000455 | hexokinase HKDC1 |
| 103818986 | MED27 | 5.91 | 0.000418 | mediator complex subunit 27 |
| 103820599 | EZR | 6.09 | 0.000584 | ezrin |
| 103823391 | ACBD6 | 6.18 | 0.000654 | acyl-CoA binding domain containing 6 |
| 103816299 |  | 9.94 | 0.000382 | protein eva-1 homolog C-like |
| 103813627 | FAM234B | 21.5 | 0.000378 | family with sequence similarity 234 member B |
| 103824821 | INSRR | 31.2 | 0.000383 | insulin receptor related receptor |
